# Subunit-specific roles of LRRC8 proteins in determining glutamate permeability of astrocytic volume-regulated anion channels

**DOI:** 10.1101/2025.07.24.666608

**Authors:** Madison L. Chandler, Ariella D. Sprague, Julia W. Nalwalk, Alexander A. Mongin

## Abstract

Volume-regulated anion channels (VRACs) are ubiquitous chloride channels that play important roles in cell volume regulation and numerous other physiological processes. VRACs are heteromeric complexes composed of leucine-rich repeat-containing proteins LRRC8A-E. LRRC8 subunit composition determines biophysical properties of VRACs, including permeability to small signaling molecules. Here, we used primary astrocyte cultures from wild-type and genetically modified C57BL/6 mice to investigate (i) subunit composition of native VRACs in the brain and (ii) subunit determinants of VRAC permeability to the excitatory neurotransmitter glutamate. qRT-PCR and RNA-seq analyses revealed high expression of *Lrrc8a-d* in mouse forebrain and astrocytes. Genetic deletion of the essential LRRC8A protein abolished swelling-activated glutamate release, measured as efflux of the non-metabolizable D-[^3^H]aspartate, confirming the crucial role of VRACs in this process. RNAi-mediated knockdown of individual subunits identified LRRC8A and LRRC8C as key components of glutamate-permeable astrocytic VRACs. qRT-PCR and Western blot analyses further showed that knockdown of LRRC8A or LRRC8C reciprocally altered the protein stability of the partner subunit without affecting their mRNA levels. A similar pattern of mutual regulation was observed between LRRC8A and LRRC8D. In contrast to LRRC8C, downregulation of LRRC8D had a more limited impact on glutamate release. Additional double-knockdown experiments demonstrated that LRRC8C- and LRRC8D-containing channels form distinct VRAC populations. This model was further supported by Western blot results showing no reciprocal regulation of LRRC8C and LRRC8D stability. Together, these findings refine our understanding of how the subunit organization of native brain VRACs governs gliotransmitter release, with implications for normal brain function and neurological disease.

**NEW & NOTEWORTHY:** Volume-regulated anion channels (VRACs) are ubiquitously expressed chloride channels composed of LRRC8A-E proteins. In human disorders, gain or loss of VRAC function leads to severe neurological phenotypes, potentially due to altered release of amino acid neurotransmitters. Here, we show that glutamate-permeable VRACs in brain astrocytes are primarily composed of LRRC8A and LRRC8C proteins. These findings provide insight into subunit organization of native VRACs in the CNS, with implications for normal brain function and neurological disease.

## INTRODUCTION

Volume-regulated anion channels (VRACs), also known as volume-sensitive outwardly rectifying anion channels (VSOR) or volume-sensitive organic osmolyte/anion channels (VSOAC), are swelling-activated chloride (Cl⁻) channels found in virtually all vertebrate cells (1-3). VRAC activity is typically assessed by measuring swelling-induced Cl⁻ currents (I_Cl,vol_) in cells exposed to hypoosmotic media [e.g., early studies (4-7)]. However, in addition to Cl⁻, VRACs are also permeable to a range of small negatively charged or uncharged organic molecules, collectively referred to as organic osmolytes (1, 8-11). As their name implies, VRACs play a key role in cell volume control and mediate regulatory volume decrease (RVD), a fundamental physiological process that restores cell volume after osmotic swelling. RVD is driven primarily by the electrically coupled efflux of cytosolic Cl⁻ (via VRACs) and K⁺ (via separate K^+^ channels), with additional contributions from bicarbonate and organic osmolytes (12, 13). Beyond their role in RVD, VRACs have been implicated in numerous other physiological and pathological processes, including regulation of membrane potential, cell proliferation, apoptosis, immune responses, and paracrine signaling [reviewed in (14-19)].

Historically, investigation of cell- and tissue-specific VRAC functions was hindered by the unknown molecular identity of these channels (20-22). A major breakthrough occurred in 2014 when the laboratories of Ardem Patapoutian and Thomas Jentsch independently used genome-wide siRNA screens to identify leucine-rich repeat-containing family 8 member A (LRRC8A, also known as SWELL1) as an essential component of VRACs (23, 24). LRRC8A belongs to a family of five LRRC8 proteins (LRRC8A-E) that share an evolutionary origin with vertebrate pannexins (25). Whole-body deletion of LRRC8A is embryonically and postnatally lethal, with ∼80% of LRRC8A-null mice dying in utero and the remaining live-born animals succumbing rapidly to multi-organ failure (26). In contrast, tissue- and cell type-specific deletions of LRRC8A are generally better tolerated [e.g., (27-29)]. Consistent with this pattern, mice with brain-specific LRRC8A knockout (bLRRC8A KO, generated by Nestin^Cre^-driven recombination) are born at expected Mendelian ratios and display no overt abnormalities in brain development (30). However, bLRRC8A KO animals unexpectedly develop spontaneous seizures and all die during late adolescence or early adulthood (30, 31), highlighting the critical yet poorly understood role of VRACs in the regulation of neural excitability.

Although VRACs can modulate neuronal activity via multiple mechanisms, their influence over brain excitability is usually attributed to their permeability to amino acid-derived neurotransmitters and modulators (15, 32-34). VRACs conduct excitatory glutamate and aspartate, as well as inhibitory GABA and taurine (8, 9, 11). Within the brain, astrocytes are thought to be the main site of physiological and pathological VRAC activity due to the propensity of these cells to swell in response to neuronal excitation or brain injury (34-36). Under physiological conditions, limited opening of astrocytic VRACs is also seen in response to GPCR-driven increases in intracellular Ca^2+^ (28, 37-39). In vivo, conditional deletion of astrocytic LRRC8A reduces tonic glutamate and GABA signaling and modifies animal behavior (28, 40). VRACs’ influence becomes even more pronounced in neural pathologies that promote pathological cell swelling, such as stroke, traumatic brain injury, and epilepsy (15, 34-36). In animal stroke models, putative but structurally diverse VRAC blockers—tamoxifen, DCPIB, dicoumarol, and bromadiolone—strongly reduce ischemic brain injury (41-45), and this effect is partially recapitulated in bLRRC8A KO mice (28, 31, 46), highlighting VRAC as an attractive therapeutic target.

Given the emerging importance of VRACs in brain health and disease, there is strong interest in their molecular structure and function. Cryo-electron microscopy studies have established that LRRC8 proteins assemble into hexameric channels with one central ion-conducting pore (47-52). The essential LRRC8A protein must co-assemble with at least one additional isoform—LRRC8C, LRRC8D, or LRRC8E—to form fully functional VRACs (24). The partnering LRRC8B-E subunits are expressed in a cell- and tissue-specific manner and confer functional diversity to VRACs, including differential permeability to organic osmolytes (16, 18, 19, 21). For example, LRRC8A/LRRC8D heteromers preferentially conduct small uncharged molecules and zwitterions, including myo-inositol, taurine, GABA, and glutamine (53-55). In contrast, there is little consensus on the subunit organization of VRACs permeable to negatively charged glutamate and aspartate, with several studies reporting different LRRC8 assemblies depending on cell type [see refs. (54-56) and Discussion]. This represents an important gap in knowledge since VRAC-driven glutamate release is thought to play an important role in both the physiological and pathological regulation of brain excitability. Hence, in the present study, we sought to identify the LRRC8 subunits that confer glutamate permeability to endogenous VRACs in primary mouse astrocytes.

## MATERIALS AND METHODS

### Ethical Approval

All animal procedures used in the present study were approved by the Institutional Animal Care and Use Committee of Albany Medical College (ACUP 24-03002), and strictly conformed to the Guide for the Care and Use of Laboratory Animals as adopted by the U.S. National Institutes of Health (https://grants.nih.gov/grants/olaw/guide-for-the-care-and-use-of-laboratory-animals.pdf).

### Animals

This study utilized two types of mice to collect brain tissue and generate primary astrocyte cultures. Wild type (WT) animals were of the C57BL/6J background, originally obtained from the Jackson Laboratory (Farmington, CT, USA; strain #000664, RRID: IMSR_JAX:000664) and bred in-house. Additionally, we used mice with conditional knockout of LRRC8A in the brain (bLRRC8A KO). These were produced by breeding *Lrrc8a*^flox/flox^ females (gift of Dr. Rajan Sah, Washington University School of Medicine, St. Louis, MO, USA) with commercially available *Nestin*^Cre/+^ males (B6.Cg-Tg(Nes-cre)1Kln/J; Jackson Lab stock #003771, RRID:IMSR_JAX:003771). Heterozygous *Nestin*^Cre/+^:*Lrrc8a*^flox/+^ males were further crossed with *Lrrc8a*^flox/flox^ female mice to produce *Nestin*^Cre/+^:*Lrrc8a*^flox/flox^ mice (bLRRC8A KO) and littermate controls. The specificity and efficacy of this genetic strategy were thoroughly validated in our previous work (30, 46). To visualize Cre activity and monitor for unintentional germline recombination, mice were additionally inbred with a strain carrying the Ai9 tdTomato Cre reporter (B6.Cg-Gt(ROSA)26Sor^tm9(CAG-tdTomato)Hze^/J; Jackson Lab stock #007909, RRID:IMSR_JAX:007909). For simplicity, the presence of the reporter allele is not shown in the breeding diagram. All mice were housed in a temperature, humidity, and light-controlled facility on a 12:12 h light/dark cycle and given free access to food and water.

### Brain Tissue Collection and Processing

For brain tissue collection, 14-week-old WT mice were euthanized with a lethal injection of sodium pentobarbital and perfused transcardially, first with room temperature phosphate-buffered saline (PBS), followed by ice-cold PBS. For qRT-PCR analysis, brains were rapidly removed and immediately transferred to a large dissection dish containing ice cold PBS additionally treated with the RNase inhibitor diethyl pyrocarbonate (DEPC, 0.1%). The olfactory bulbs, cerebellum and brainstem were removed, and a midline sagittal cut was made to separate the hemispheres. Each hemisphere was snap-frozen in liquid nitrogen and stored at –80°C until further processing.

### Primary Cultures of Mouse Astrocytes

Primary cultures of mouse astrocytes were prepared from brain cortices of newborn (P0-P2) pups of both sexes as most recently described in (46) with minor modifications. Briefly, neonates were anesthetized by cooling and rapidly decapitated. Brains were harvested and transferred into ice-cold sterile Dulbecco’s PBS without calcium and magnesium (ThermoFisher Scientific, Waltham, MA, USA, cat. #14190144). Cortical tissue was dissected from meninges, minced using a sterile surgical blade, incubated with 500 µL of the recombinant protease TrypLE Express (ThermoFisher, cat. #12605) and additionally triturated by pipetting. Triturated tissue was digested with TrypLE for 5 min at room temperature (22°C). Enzymatic digestion was terminated by adding Earl’s minimal essential medium (MEM, ThermoFisher, cat. #11095) supplemented with 10% heat-inactivated horse serum (HIHS, ThermoFisher, cat. #26050) and penicillin/streptomycin (P/S, ThermoFisher, cat. #15140). The cell suspension was next filtered through a 70-µm nylon mesh cell strainer (Corning, Corning, NY, USA, cat. # 431751). To obtain an enriched astrocyte population, cells extracted from either half brain (for WT), or full brain (bLRRC8A KO) were separately plated onto T75 flasks and grown in a humidified atmosphere of 5% CO_2_/balance air at 37°C. With this approach, primary cell cultures typically reached confluency after around two weeks and were maintained for up to six weeks until replating for functional assays. The purity of astrocyte populations was periodically verified with immunofluorescence by staining for the astrocyte marker, glial fibrillary acidic protein (anti-GFAP, MilliporeSigma, St. Louis, MO, USA, cat. #MAB360, RRID:AB_11212597; 1:400 dilution) and was ≥95%.

### Quantitative Real-Time PCR (qRT-PCR)

Relative gene expression and efficacy of siRNA gene knockdowns were assessed using qRT-PCR. Total RNA was isolated from brain tissue or primary astrocyte cultures using the Aurum Total RNA Mini Kit (Bio-Rad, Hercules, CA, USA, cat. #732-6820) following the manufacturer’s instructions. Processing of tissue samples included on-column DNase digestion to eliminate genomic DNA contamination. RNA concentration was measured with a NanoDrop 1000 spectrophotometer (ThermoFisher, RRID:SCR_016517). RNA was immediately converted to cDNA using the iScript cDNA synthesis kit (Bio-Rad, cat. #1708891), with 1 µg of total RNA per 20 µL reaction. The resulting cDNA was analyzed for the relative abundance of *Lrrc8* transcripts by qRT-PCR using gene-specific primers and SYBR Green Supermix (Bio-Rad, cat. #1708882) in a CFX96 Real-Time PCR Detection System (Bio-Rad, RRID:SCR_018064). Expression levels were normalized to the housekeeping gene *Rpl13a*, and where applicable, further normalized to control samples using the 2^-ΔΔC(T)^ method (57). As an internal quality control, expression of an additional housekeeping gene, *Rps20*, and the abundant transcript *Gapdh* were also quantified. All qRT-PCR primers were obtained as validated GeneGlobe assays from Qiagen (Germantown, MD, USA); detailed catalog information is provided in Table 1.

**Table 1.** qRT-PCR primers used for gene expression analysis.

| Primer target | Primer ID | Source | GeneGlobe ID |
| --- | --- | --- | --- |
| *Rpl13a* (housekeeping) | Mm_Rpl13a_1_SG | Qiagen | QT00267197 |
| *Rps20* (housekeeping) | Mm_Rps20_1_SG | Qiagen | QT00251433 |
| *Gapdh* (housekeeping) | Mm_Gapdh_3_SG | Qiagen | QT01658692 |
| *Lrrc8a* | Mm_Lrrc8a_2_SG | Qiagen | QT01167502 |
| *Lrrc8b* | Mm_Lrrc8b_1_SG | Qiagen | QT01541666 |
| *Lrrc8c* | Mm_Lrrc8c_1_SG | Qiagen | QT01042881 |
| *Lrrc8d* | Mm_Lrrc8d_1_SG | Qiagen | QT02255085 |
| *Lrrc8e* | Mm_Lrrc8e_1_SG | Qiagen | QT01046269 |

### Western Blotting

Protein levels of LRRC8A, LRRC8C, and LRRC8D were measured in lysates from primary astrocyte cultures. Cells were lysed in a denaturing buffer containing 2% sodium dodecyl sulfate (SDS) and 8 mM EDTA. Protein concentration was determined using the Pierce Bicinchoninic acid protein assay kit (Thermo Fisher Scientific, cat. #23225). The remaining lysate was diluted with 4× Laemmli sample buffer (Bio-Rad, cat. #4561033) and stored at −80^◦^C until analysis. Lysates were boiled for 5 min, and 10-20 µg of total protein per sample was resolved on 10% Mini-PROTEAN TGX gels (Bio-Rad, cat. #1610747) via SDS-PAGE. Proteins were then transferred to polyvinylidene difluoride (PVDF) membranes (Bio-Rad, cat. #1620177) using a Trans-Blot Turbo transfer system (Bio-Rad, RRID:SCR_023156). Membranes were blocked for 20 min in Tris-buffered saline with 0.1% Tween-20 (TBS-T) containing 5% non-fat dry milk and further incubated with primary antibody overnight at 4^◦^C. After washing in TBS-T, membranes were incubated for 2 h at room temperature with a species-matching HRP-conjugated secondary antibody in 5% milk/TBS-T. Protein bands were visualized using enhanced chemiluminescence reagent (ECL; GE Healthcare, cat. #RPN2232) and a ChemiDoc Imaging System (Bio-Rad, RRID:SCR_019684). For loading controls, membranes were washed and re-probed for 30 min with HRP-conjugated anti-β-actin antibody. Band intensities were quantified using ImageJ software (58) (NIH, RRID:SCR_003070) and were normalized to β-actin immunoluminescence from the same blot. Detailed information on all antibodies and their dilutions is provided in Table 2. For each antibody, we provide representative full-length Western blot images in the main figures. The complete dataset of analyzed Western blots, including matching β-actin loading controls, is included with Supplemental materials.

**Table 2.** Primary and secondary antibodies used for western blot analysis.

| Protein target | Species/type | Source | Catalog # | RRID | Dilution |
| --- | --- | --- | --- | --- | --- |
| β-actin, HRP-conjugated | Mouse, monoclonal | Millipore-Sigma | A3854 | AB_262011 | 1:50,000 |
| LRRC8A | Mouse monoclonal | Santa Cruz Biotech. | sc-517113 | AB_2928142 | 1:1,000 |
| LRRC8C | rabbit polyclonal | Proteintech | 21601-1-AP |  | 1:2,000 |
| LRRC8D | rabbit polyclonal | Proteintech | 11537-1-AP | AB_2137767 | 1:2,000 |
| anti-mouse IgG, HRP-conjugated | sheep | GE Healthcare | NA931 |  | 1:10,000 |
| anti-rabbit IgG, HRP-conjugated | donkey | GE Healthcare | NA934 |  | 1:10,000 |

### RNAi-Mediated LRRC8 Gene Silencing

Targeted knockdown of LRRC8 family members was achieved using gene-specific siRNA constructs obtained from Qiagen (Valencia, CA, USA). AllStars Negative Control siRNA (Qiagen) was used as a non-targeting control. Primary astrocytes at ∼70-80% confluency were transfected using Lipofectamine RNAiMAX (Thermo Fisher, cat. #13778075) according to the manufacturer’s protocol. siRNA-Lipofectamine complexes were prepared in Opti-MEM (Thermo Fisher, cat. #31985), diluted further with Opti-MEM, and added to cells at final siRNA concentrations ranging from 10 to 50 nM (optimized for individual siRNAs). Details of the siRNA constructs, including catalog numbers and target sequences, are listed in Table 3. Knockdown efficacy for each construct was validated by qRT-PCR. After a 4h incubation, cultures were supplemented with MEM containing 10% HIHS and penicillin plus streptomycin (MEM+HIHS+P/S). At 24 h post-transfection, the medium was fully replaced with fresh MEM+HIHS+P/S. mRNA expression levels were assessed 48h post-transfection, and functional assays were performed 96h post-transfection.

**Table 3.**
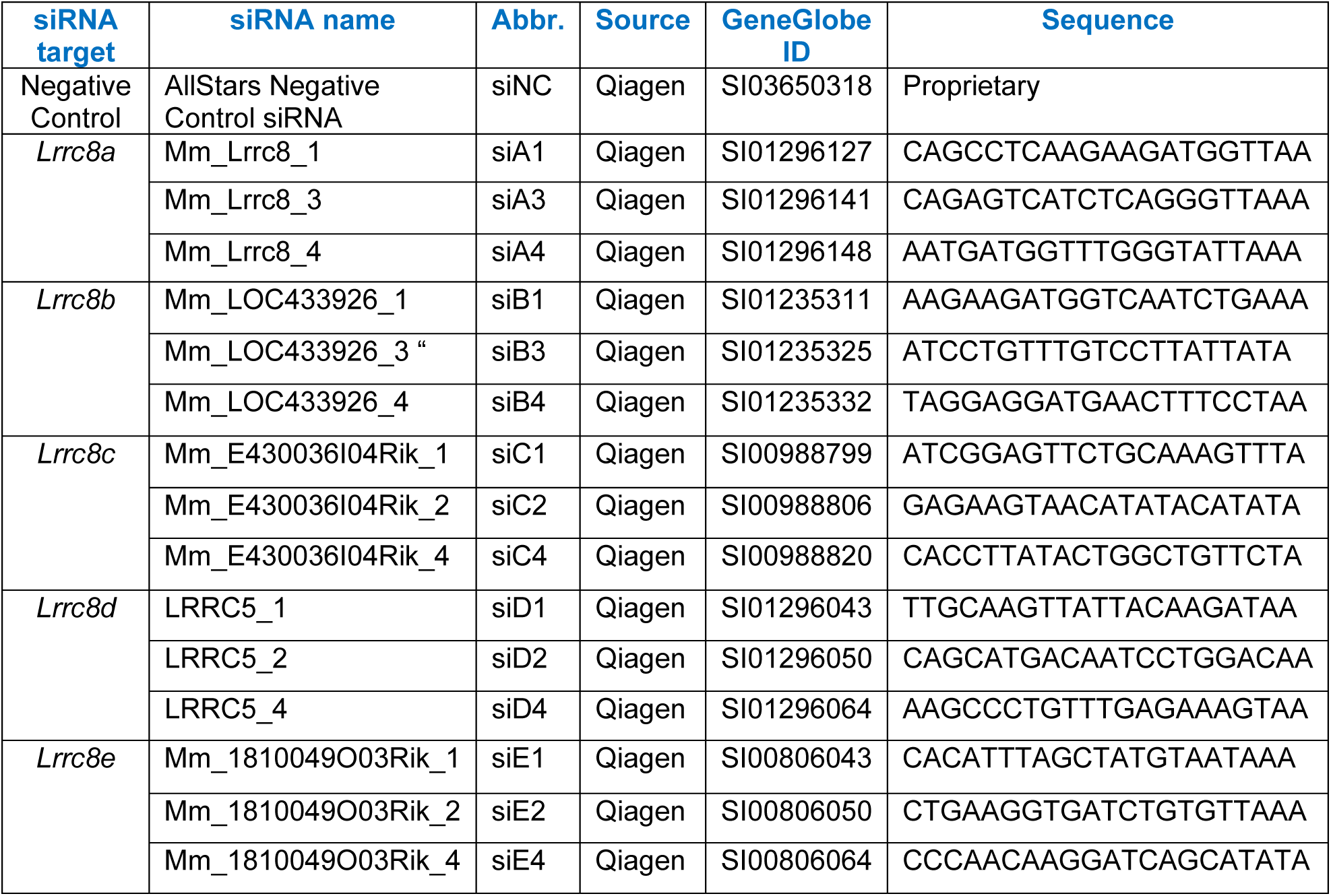
siRNA constructs used in RNAi experiments.

### Radiotracer Assays of VRAC Activity

VRAC activity was evaluated by measuring the release of D-[^3^H]aspartate, a non-metabolizable analog of glutamate, as extensively validated in our earlier studies [e.g., (55, 59, 60)]. Two complementary assay formats were used, employing astrocytes cultured either in multi-well plates or on glass coverslips. In all experiments, astrocytes were preloaded overnight in standard serum-containing medium supplemented with 2 μCi/mL D-[^3^H]aspartate (Revvity, cat. #NET50100).

For medium-throughput screening of multiple siRNA constructs, cells were grown on poly-D-lysine-coated 12-well plates. After radiotracer loading, cultures were washed to remove extracellular isotope and equilibrated for 30 min at 37°C in isoosmotic Basal medium containing (in mM): 135 NaCl, 3.8 KCl, 1.2 MgSO_4_, 1.3 CaCl_2_, 1.2 KH_2_PO_4_, 10 D-glucose, and 10 HEPES (pH 7.4; osmolality 290 ±3 mOsm/kg). To inhibit radiotracer reuptake via Na⁺-dependent GLAST transporters, cells were exposed to LiCl-substituted media, in which NaCl was isoosmotically replaced with LiCl (61). Baseline release was measured during a 10 min incubation in isoosmotic Li⁺-containing medium, followed by stimulation with hypoosmotic Li⁺ medium (LiCl reduced by 50 mM; final osmolality ∼200 mOsm/kg, ∼30% reduction) to activate VRACs. Ten-min fractions of isoosmotic and hypoosmotic media were collected for analysis. Radiotracer release was quantified by liquid scintillation counting using a TRI-CARB 4910TR counter (Revvity), after mixing samples with Ecoscint A scintillation fluid (National Diagnostics, cat. #LS-273). Integral release values were normalized to the remaining intracellular D-[^3^H]aspartate, determined after cell lysis in 2% SDS containing 8 mM EDTA.

To resolve the kinetics of VRAC activation, D-[^3^H]aspartate efflux assays were conducted using a custom-fabricated Lucite superfusion chamber that has a depression on the bottom to accommodate an 18 × 18 mm glass coverslip and a Teflon screw lid that leaves a ∼200 μm space above the cells. Coverslips with D-[^3^H]aspartate-preloaded astrocytes were briefly washed with Basal medium (for composition see above), transferred into the chamber, and superfused at a constant rate of ∼1.4 mL/min with either isoosmotic or hypoosmotic media. In this setup, we used NaCl-containing media because released D-[^3^H]aspartate was effectively removed with the flow. One-min perfusate fractions were collected into scintillation vials using an automated CF-1 fraction collector (SpectraLab Scientific, Markham, ON, Canada). At the conclusion of each experiment, cells were lysed in 2% SDS+8 mM EDTA, and D-[^3^H]aspartate content in all samples was quantified using a scintillation counter as described above. Data were expressed as fractional release values, calculated at each time point as a ratio of released [^3^H] to the total [^3^H] content (cumulative release + remaining tracer), using a custom Excel-based analysis program.

### Statistics

All data are presented as mean values ± SD. The n value corresponds to an individual animal (qRT-PCR and RNA-seq) or an independently transfected astrocyte culture. For cell culture experiments, all results were reproduced in multiple cultures prepared from at least two different animals, unless specified otherwise. Statistical differences between three or more groups were analyzed using one-way ANOVA followed by Tukey’s multiple comparisons test. In high-throughput functional assays, we used repeated-measures two-way ANOVA to assess the effects of medium (cell swelling), treatment (siRNA), and their interaction, followed by Tukey’s multiple comparisons test. In several instances, to increase the clarity of the graphical presentation, we show only the most germane statistical effects; full analyses are included in the primary datasets deposited in the Open Science Framework (https://osf.io/xvyba). In RNAi expression assays, mRNA expression values were normalized to the expression levels in siNC-treated cells and compared to unity using a one-sample t-test with Bonferroni correction for multiple comparisons. All statistical analyses and graphical preparations were performed using GraphPad Prism 10.0 (GraphPad Software, San Diego, CA, USA; RRID:SCR_002798).

## RESULTS

### LRRC8A Deletion Abolishes Activity of Glutamate-Permeable VRACs in Astrocytes

To investigate the role of LRRC8 proteins in the formation of endogenous astrocytic VRACs, we first employed a molecular genetics approach using *Nestin*^Cre^-driven recombination. *Nestin*^Cre^ targets neural precursor cells, leading to gene deletion in all cells of neuroectodermal origin, including astrocytes (62, 63). Our conditional knockout breeding strategy produced four genotypes: two types of controls (*Lrrc8a*^flox/+^ and *Lrrc8a*^flox/flox^ without Cre, referred to as fl/+ and fl/fl), a heterozygous (HET) LRRC8A deletion, and a homozygous LRRC8A knockout (KO) (Fig. 1A). Primary astrocyte cultures were prepared from neonatal brains of each genotype and successful Cre recombination was confirmed using the Ai9 tdTomato reporter. Live-cell imaging of tdTomato fluorescence showed complete Cre penetrance in both LRRC8A HET and KO astrocytes (representative images in Fig. 1B). Western blot analysis of astrocyte culture lysates revealed that heterozygous LRRC8A deletion reduced LRRC8A protein levels by approximately 60% (p < 0.0001, Fig. 1D), while KO cultures showed a complete loss of LRRC8A immunoreactivity (p < 0.0001, Fig. 1D, full western blot dataset in Supplemental Fig. S1).

**Figure 1:**
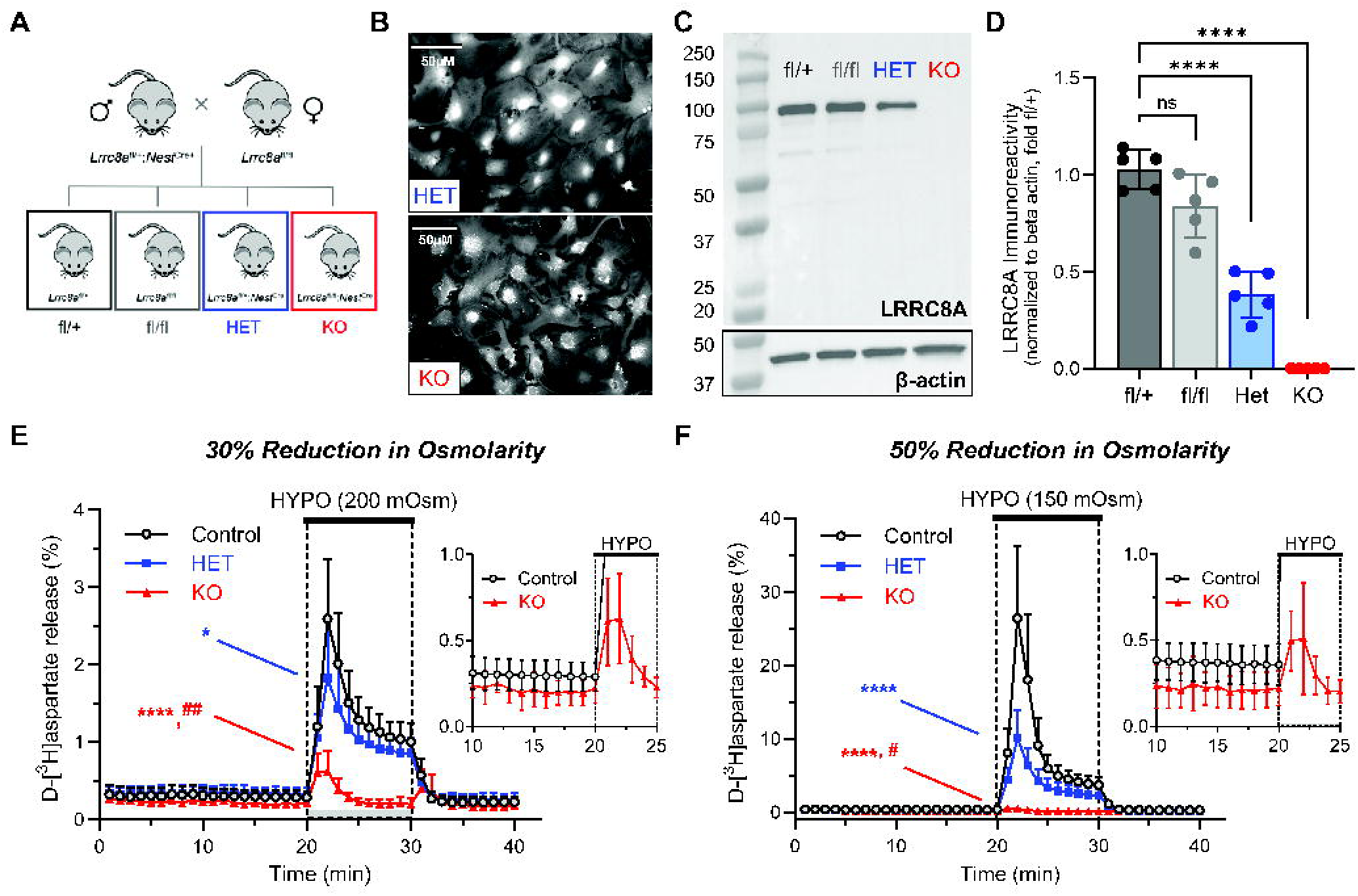
LRRC8A-containing VRACs are the main conduit for swelling-activated release of cytosolic glutamate. **A**, Breeding strategy for brain-specific deletion of LRRC8A using *Nestin*^Cre^ recombination. The four genotypes generated are: *Lrrc8a*^flox/+^ (fl/+), *Lrrc8a*^flox/flox^ (fl/fl), *Lrrc8a*^flox/+^:*Nestin*^Cre/+^ (HET), and *Lrrc8a*^flox/flox^:*Nestin*^Cre/+^ (KO). **B**, Representative live images of LRRC8A HET and KO astrocytes expressing the Cre reporter Ai9. TdTomato fluorescence was captured at 320× magnification. **C**, Representative full-length Western blot of LRRC8A immunoreactivity and matching β-actin signal in primary astrocyte cultures from the four genotypes shown in (A). **D**, Quantification of LRRC8A expression normalized to β-actin and presented relative to the average signal from fl/+ samples. Data are the mean values ± SD from five cultures per group. ****p < 0.0001, one-way ANOVA with Tukey’s multiple comparisons test. **E**, VRAC activity in primary astrocytes assessed as swelling-activated release of D-[^3^H]aspartate in response to a 30% reduction in medium osmolality (290 to 200 mOsm). Data are the mean values ± SD from 8-16 independent experiments in at least two different astrocyte cultures per genotype. *p < 0.05, ****p < 0.0001, HET (n=8) and KO (n=8) vs. controls (fl/+ and fl/fl, n=16), respectively. ^##^p < 0.01, HET vs. KO. One-way ANOVA of maximal release values with Tukey’s multiple comparisons test. **Inset**: Hypoosmotic response in LRRC8A KO astrocytes shown on an expanded scale. **F**, VRAC activity in primary astrocytes exposed to a 50% reduction in medium osmolality (from 290 to 150 mOsm). Data are the mean values ± SD from 8–11 independent experiments in at least two different astrocyte cultures per genotype. ****p < 0.0001, HET (n=9) and KO (n=8) vs. controls (fl/+ and fl/fl, n=11); ^#^p < 0.05, HET vs. KO. One-way ANOVA with Tukey’s multiple comparisons test. **Inset**: Hypoosmotic response in LRRC8A KO astrocytes shown on an expanded scale.

We next assessed functional VRAC activity using a D-[^3^H]aspartate efflux assay. In control astrocytes, a 30% reduction in extracellular osmolality (to 200 mOsm) triggered a rapid increase in D-[^3^H]aspartate release, reaching a peak rate of 2.6% within two minutes (an 8.7-fold rise over baseline), followed by a gradual decline consistent with the volume regulatory process. This kinetic profile closely matched our previous measurements of VRAC activation in both rat and mouse astrocytes (55, 59). Astrocytes with heterozygous LRRC8A deletion exhibited a 30% reduction in swelling-activated radiotracer efflux (p = 0.038, Fig. 1E, full analysis in Supplemental Fig. S2). As anticipated, LRRC8A KO astrocytes displayed a profound ∼76% decrease in swelling-induced glutamate release (p < 0.0001, Fig. 1E and Supplemental Fig. S2). However, a small, transient release component persisted during the initial three minutes of hypoosmotic exposure (Fig. 1E). This residual response was characterized by more rapid activation and inactivation kinetics as compared to typical VRAC-mediated release and is likely mediated by a distinct mechanism (see additional considerations in the following paragraph and extended Discussion). Interestingly, LRRC8A KO astrocytes also showed a substantial reduction in baseline glutamate release under isoosmotic conditions: 28% lower than that of littermate controls (p < 0.0001 see inset in Fig. 1E). Together, these findings demonstrate that (i) LRRC8A containing VRACs are the primary conduit for swelling-induced glutamate release in astrocytes, and (ii) these channels also contribute to tonic glutamate release under resting, isoosmotic conditions.

To further investigate the apparent discrepancy between the 60% decrease in LRRC8A protein levels and the relatively modest 30% inhibition of swelling-activated glutamate release in HET cells, we performed additional functional assays using media with a more pronounced reduction in osmolality. When medium osmolality was decreased by ∼50% (to 150 mOsm), VRAC activation was dramatically enhanced, compared to the response previously observed with a 30% (to 200mOsm) osmotic reduction, resulting in an over 10-fold increase in maximal release rates between the two conditions (compare Y-axis scales in Fig. 1E and 1F). LRRC8A KO cells again showed a significant reduction in baseline glutamate release, a 40% decrease as compared to controls (p < 0.0001, Fig. 1F, consistent with Fig. 1E). Importantly, in severely swollen astrocytes, heterozygous LRRC8A deletion caused a much more pronounced 62% reduction in VRAC-mediated activity, which is close to what would be expected based on changes in protein expression in HET cells (p < 0.0001, Fig. 1F, full analysis in Supplemental Fig. S3). These results may suggest that a reserve pool of LRRC8A-containing anion channels in the HET genotype can compensate under moderate swelling conditions, but this compensatory capacity is lost with more extreme osmotic stress/cellular swelling (see Discussion). This finding has implications for interpreting subsequent RNAi-based knockdown experiments.

Finally, while comparing the “residual” D-[^3^H]aspartate release in LRRC8A KO astrocytes exposed to 30% and 50% reductions in medium osmolality, we found that the magnitude of this release was indistinguishable between the two hypoosmotic conditions (maximal release rates: 0.62 ± 0.25% vs. 0.51 ± 0.32%, p = 0.498; insets in Fig. 1E and 1F). This was in stark contrast to control astrocytes, in which VRAC activity increased by an order of magnitude under stronger osmotic stress (2.59 ± 0.55% vs. 26.38 ± 5.19%, p < 0.0001). This lack of sensitivity to the extent of cellular swelling further supports the conclusion that the residual release observed in LRRC8A KO astrocytes is not mediated by VRAC (see Discussion). To explore alternative mechanism(s), specifically, the contribution of Ca^2+^-activated anion channels, we stimulated control and LRRC8A KO astrocytes with the P2Y receptor agonist ATP (64). In these experiments, ATP triggered a small, transient glutamate release with rapid inactivation kinetics (Supplemental Fig. S4). This response partially resembled the “residual” glutamate release observed in LRRC8A KO astrocytes under hypoosmotic conditions. Nevertheless, these results were inconclusive due to the small amplitude and high variability of the ATP effects in both cell types.

### Relative Abundance of *Lrrc8* Transcripts in Mouse Brain and Primary Astrocyte Cultures

We next measured and compared the endogenous expression of all five *Lrrc8* isoforms in C57BL/6J mouse brains and primary astrocyte cultures derived from C57BL/6J neonates. qRT-PCR analysis of samples prepared from forebrain tissue detected high levels of four *Lrrc8* transcripts, *Lrrc8a-Lrrc8d*, with the order of abundance *Lrrc8b>Lrrc8a≈Lrrc8d>Lrrc8c* (Fig. 2A). mRNA levels of the *Lrrc8* subunits were ∼1-5% of the housekeeping gene ribosomal *Rpl13a*, or 0.5-2.5% of the highly prevalent *Gapdh* (Fig. 2A and Supplemental Fig. S5A). Consistent with literature reports, brain levels of *Lrrc8e* were very low-to-non-detectable (Fig. 2A). To independently validate our findings on *Lrrc8* expression, we additionally performed an in-silico analysis of publicly available RNAseq datasets. Fig. 2B summarizes expression levels of the five *Lrrc8* isoforms in seven forebrain regions from five mice of both sexes from the Human Protein Atlas. The pattern of relative abundance of *Lrrc8* subunits in RNAseq datasets was nearly identical to that found in our qRT-PCR experiments: *Lrrc8b>Lrrc8d≈Lrrc8a>Lrrc8c*, and extremely low-to-nondetectable *Lrrc8e* (compare Fig. 2A and 2B).

**Figure 2:**
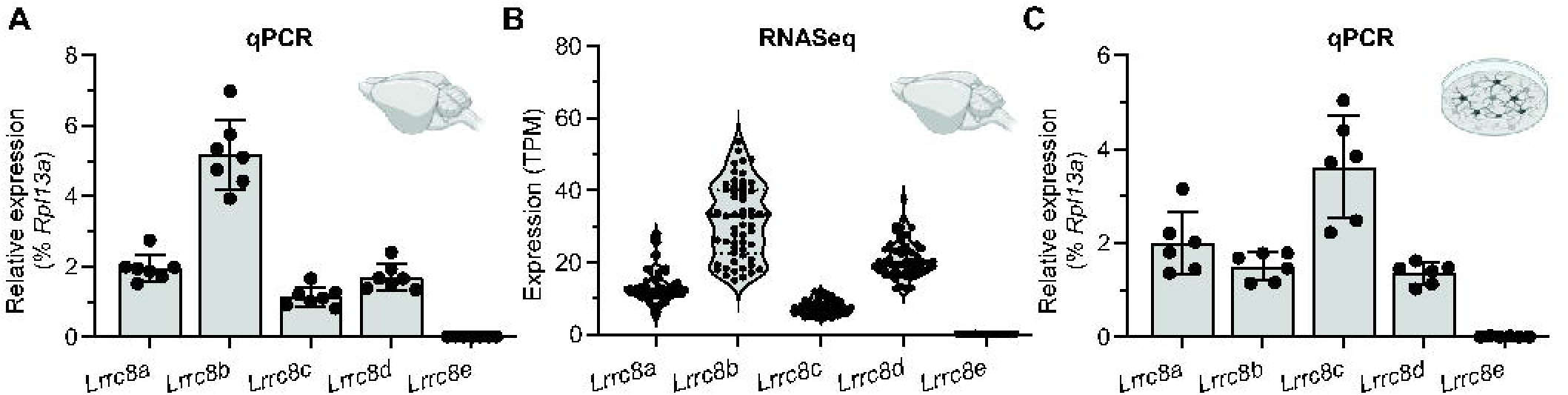
Relative abundance of *Lrrc8a-e* transcripts in mouse brain and primary astrocyte cultures. **A**, Quantification of mRNA expression levels for *Lrrc8a-e* in mouse forebrain lysates using qRT-PCR. Expression levels were normalized to ribosomal *Rpl13a* within the same sample. Data are the mean values ± SD of seven mouse forebrains. **B**, RNA-seq analysis of *Lrrc8a-e* expression in seven mouse forebrain regions from five mice of both sexes (data downloaded from the Human Protein Atlas). **C**, Quantification of mRNA expression levels for *Lrrc8a-e* in primary astrocyte cultures measured with qRT-PCR and normalized to ribosomal *Rpl13a*. Data are the mean values ± SD of six independent cultures.

For comparative purposes, we additionally analyzed *LRRC8* subunit expression in the human brain using RNAseq datasets from the Genotype-Tissue Expression (GTEx) portal [accessed on July 14, 2025]. To explore regional heterogeneity, we extracted expression data for four forebrain regions: cortex, amygdala, caudate, and hippocampus (181-300 males and females per region, Supplemental Fig. S6). Surprisingly, human tissue exhibited a strikingly different *LRRC8* expression profile as compared to the mouse brain. In humans, *LRRC8A* dominated all other LRRC8 subunits with up to 5-fold higher expression as compared to the next most abundant isoform (Supplemental Fig. S6). The relative expression levels of human VRAC subunits were *LRRC8A*>>*LRRC8B≈LRRC8D>>LRRC8C* (Supplemental Fig. S6, compare to mouse data in Fig. 2B). These species-specific differences should be considered when analyzing VRAC stoichiometry, organization, and function across different organisms and model systems.

Since our functional VRAC assays were performed in primary mouse astrocytes, we further measured *Lrrc8* mRNA levels primary astrocyte cultures using qRT-PCR. In astrocyte cultures, the average levels of *Lrrc8* transcripts were comparable to those observed in mouse forebrain tissue. Expression levels of the four predominant *Lrrc8* isoforms ranged from ∼1-4% relative to *Rpl13a*, or 0.5-2% when normalized to *Gapdh* (Fig. 2C and Supplemental Fig. S5B). As in brain tissue, *Lrrc8e* expression was detected at trace levels (Fig. 2C). Notably, we observed cell type-specific differences in the relative expression of individual *Lrrc8* isoforms: cultured astrocytes exhibited significantly higher levels of *Lrrc8c* and markedly lower levels of *Lrrc8b* compared to brain tissue (compare Fig. 2A and Fig. 2C). These differences may reflect an intrinsic cell type-specific *Lrrc8* expression profile or could result from artifacts introduced by *in vitro* culture conditions.

### Validation of RNAi Approach for Probing VRAC Function

To investigate the function of endogenous LRRC8 proteins in wild-type astrocytes, we employed RNA interference (RNAi) using commercially available siRNA constructs. The temporal dynamics and efficacy of siRNA-mediated knockdown were assessed by measuring LRRC8A protein levels at 24-96h post-transfection with an *Lrrc8a*-targeting siRNA, termed siA1. LRRC8A immunoreactivity declined progressively over time, with reductions of 46%, 87%, 93%, and 97% at 24, 48, 72, and 96h, respectively (Fig. 3A, full western blot dataset in Supplemental Fig. S7). Based on this time course, all subsequent functional assays were performed 96h after transfection. The kinetics of protein loss suggested an LRRC8A half-life of approximately 24 h (Fig. 3A).

**Figure 3:**
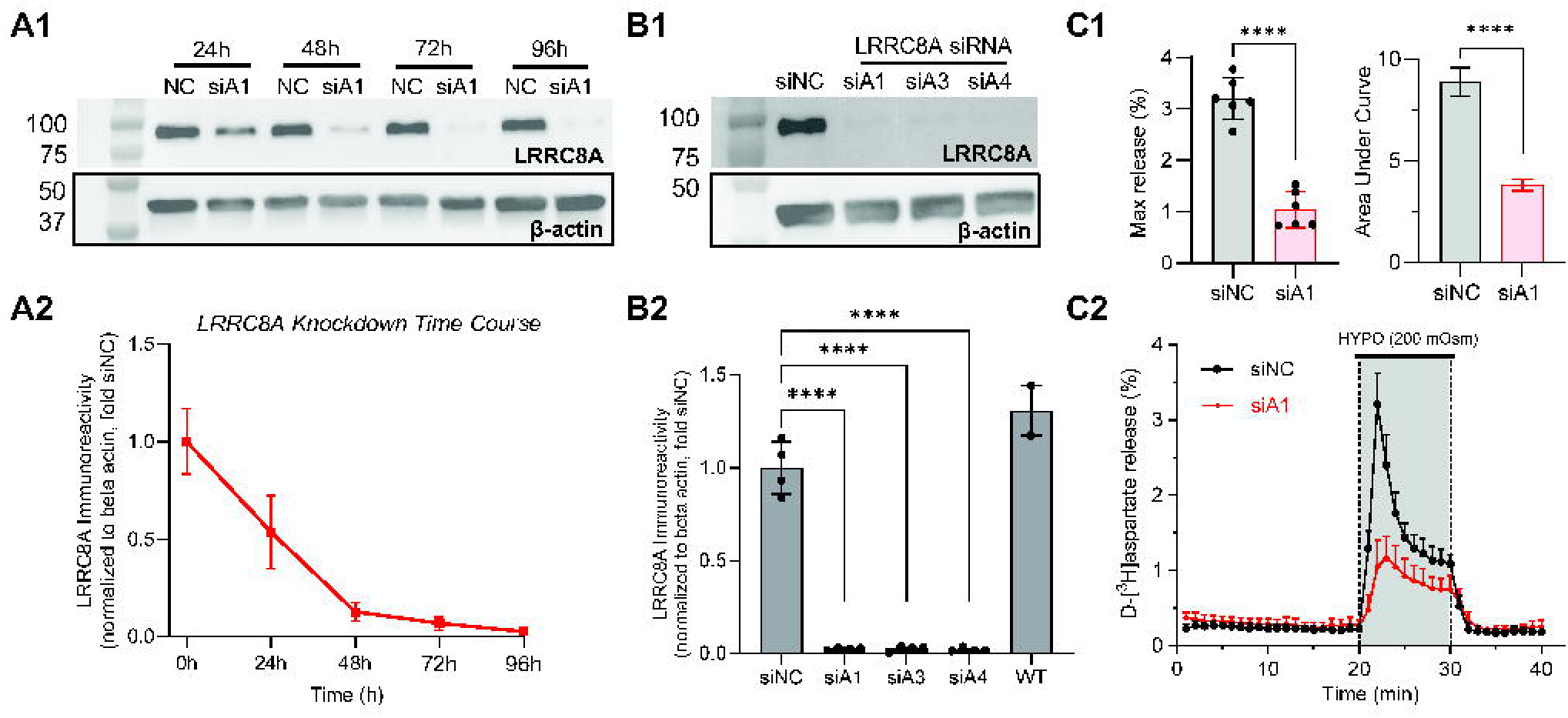
Time Course of siRNA-mediated LRRC8A knockdown and its impact on swelling-activated glutamate release. **A1**, Representative Western blot of LRRC8A immunoreactivity and corresponding β-actin signal from protein lysates collected 24h, 48h, 72h, and 96h post siRNA transfection with either negative control siRNA (NC) or *Lrrc8a-*targeting siRNA (siA1). **A2**, Quantification of LRRC8A protein levels, normalized to β-actin and presented relative to the average signal from siNC samples. Data are the mean values ± SD from three independent transfections performed in three different astrocyte cultures. **B1**, Representative Western blot of LRRC8A immunoreactivity and matching β-actin signal from protein lysates collected 96h after treatment with siNC or one of three *Lrrc8a*-targeting siRNA constructs (siA1, siA3, or siA4). **B2**, Quantification of LRRC8A protein levels normalized to β-actin and presented relative to the average signal from siNC samples. Data are the mean values ± SD from four independent transfections performed in four different astrocyte cultures. ****p < 0.0001, **p < 0.01, one-way ANOVA with Tukey’s multiple comparisons test. **C1**, Statistical comparison of the maximal D-[^3^H]aspartate release values and area under the curve from C2. Data are the mean values ± SD in six independent transfections in two different astrocyte cultures per group. ****p < 0.0001, unpaired t-test. **C2**, Kinetics of VRAC-mediated D-[^3^H]aspartate release from astrocyte cultures treated with siNC or siA1 and exposed to a 30% reduction in medium osmolality.

We next compared the efficacy of three independent *Lrrc8a*-targeting siRNAs, each directed against distinct regions of the mRNA (siA1, siA3, and siA4; see Table 3). Ninety-six hours post-transfection, all three constructs reduced LRRC8A protein levels by >90% (p < 0.0001; Fig. 3B, full western blot dataset in Supplemental Fig. S8).

To determine the functional impact of siRNA-mediated LRRC8A depletion on VRAC activity and compare it with genetic LRRC8A KO, we measured swelling-activated glutamate release using a D-[^3^H]aspartate efflux assay. Knockdown of LRRC8A significantly reduced D-[^3^H]aspartate efflux in response to a 30% reduction in medium osmolarity, with a ∼65% decrease in maximal release and ∼57% decrease in area under the curve (p < 0.0001; Fig. 3C). The magnitude of inhibition, 65% decrease, is comparable to the 76% reduction seen in LRRC8A KO astrocytes (Fig. 1E), indicating that siRNA knockdown is highly effective. The residual VRAC activity, despite near-total loss of detectable LRRC8A protein, likely reflects a non-proportional relationship between protein abundance and channel function, or limitations in the sensitivity of antibody-based detection.

### Analysis of Subunit-Specific Contributions of LRRC8B-E to Glutamate-Permeable VRACs

To assess the individual contributions of all five LRRC8 isoforms to glutamate permeability, we used an RNAi-based screening strategy. For each of the five *Lrrc8* mRNA species, we tested three distinct siRNA constructs targeting different regions of the mRNA (see Table 3 for target sequences). In cells treated with a non-targeting control siRNA (siNC), the relative expression profile of *Lrrc8* transcripts remained unchanged (compare Fig. 2C and Fig. 4A1). All tested siRNA constructs achieved 70-90% knockdown of their respective targets (Fig. 4B1-F1). Therefore, all siRNA constructs were included in the initial functional screenings.

**Figure 4:**
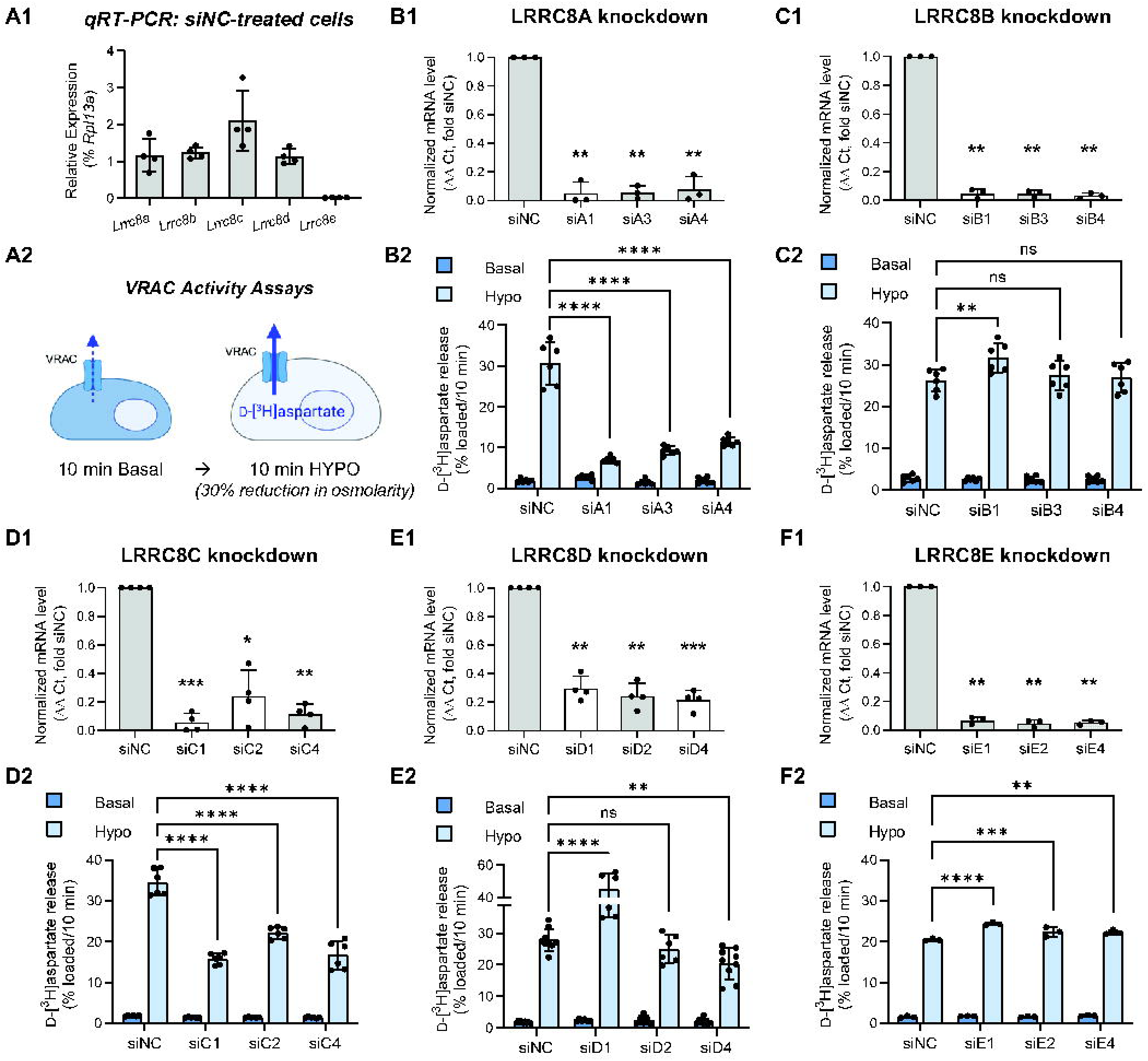
Functional contribution of individual LRRC8A-E subunits to glutamate-permeable VRACs. **A1**, Quantification of *Lrrc8a-e* mRNA expression levels in mouse astrocyte cultures treated with negative control siRNA (siNC). Expression levels were normalized to the housekeeping *Rpl13a*. Data are the mean values ± SD of four independently prepared cell cultures. **A2**, Schematic of a radiotracer assay of VRAC activity (created with BioRender.com, see Methods for additional details). **B-F**, Effects of gene-specific siRNA constructs on *Lrrc8a-e* mRNA expression levels 48 h post-transfection and swelling-activated D-[^3^H]aspartate release 96 h post transfection. ***B1-F1***, Relative *Lrrc8a-e* expression was normalized to *Rpl13a* and compared to siNC. Values are the means ± SD of 3-4 independent transfections in at least two different astrocyte cultures. ***p < 0.001, **p < 0.01, *p < 0.05, one-population t-test with Bonferroni correction for multiple comparisons. ***B2-F2***, Integral D-[^3^H]aspartate release values in cells exposed for 10 min to either isoosmotic (Basal) or hypoosmotic (Hypo) conditions (see Methods). With the exception of *F2*, data are the mean values ± SD of 6-9 independent transfections in two independently prepared cell cultures. For *Lrrc8e* knockdowns in *F2*, data are the mean values ± SD of three independent transfections in one astrocyte culture. ****p < 0.0001, *** p < 0.001, ** p < 0.01 vs. siNC, repeated measures two-way ANOVA with Tukey’s multiple comparisons test (for clarity only the effect of siRNA under hypoosmotic conditions are shown).

To increase experimental throughput, initial VRAC activity assays were conducted in a multi-well plate format, measuring swelling-activated D-[^3^H]aspartate release in response to a 30% reduction in medium osmolarity (see schematic Fig. 4A2). As detailed in the Methods section, these efflux assays were performed in Na^+^-free medium to prevent reuptake of the tracer by Na^+^-dependent GLAST transporters. For all experiments, repeated-measures two-way ANOVA revealed a main effect of medium osmolarity (basal vs. hypoosmotic; p < 0.0001), consistent with VRAC activation (Fig. 4B2-F2). For simplicity, we present only post-hoc analyses comparing the effects of individual siRNA constructs with those of siNC-treated controls under hypoosmotic conditions. We first tested the effect of siRNA-mediated knockdown of the essential LRRC8A subunit. All three siRNAs targeting LRRC8A (designated siA1, siA3, and siA4) strongly reduced swelling-induced D-[^3^H]aspartate release, decreasing it by 67-85% as compared to siNC controls (p < 0.0001 vs. siNC; Fig. 4B2). The degree of inhibition ranked siA1 > siA3 > siA4 (Fig. 4B2). Importantly, the high inhibitory potency of the LRRC8A knockdown resembled the near-complete loss of VRAC activity in LRRC8A KO astrocytes.

In contrast to the strong effect of LRRC8A knockdown, none of the siRNAs targeting LRRC8B (siB1, siB3, and siB4) inhibited VRAC activity (Fig. 4C2). However, repeated measures two-way ANOVA revealed a significant interaction between siRNA treatments and media osmolarity (p = 0.02; Fig. 4C2), so we performed a post hoc analysis. Interestingly, one construct, siB1, produced a statistically significant 24% increase in swelling-activated D-[^3^H]aspartate release (p = 0.002 vs. siNC; Fig. 4C2). For all subsequent experiments, we used siB4, which demonstrated high efficacy in reducing *Lrrc8b* mRNA levels, without affecting functional VRAC activity.

All three siRNA constructs targeting LRRC8C (siC1, siC2, and siC4) significantly reduced VRAC activity (p < 0.0001 vs. siNC; Fig. 4D2). The most effective, siC1, reduced swelling-activated D-[^3^H]aspartate release by 56% (p < 0.0001 vs. siNC; Fig. 4D2). The order of inhibition was siC1 > siC4 > siC2, which closely mirrored the efficacy of *Lrrc8c* mRNA knockdown (Fig. 4D1). Due to its superior efficacy, siC1 was used in all subsequent experiments. Western blotting confirmed effective LRRC8C protein depletion at the time of functional analysis, with a ∼79% reduction in LRRC8C immunoreactivity 96h after siRNA transfection (p < 0.0001; Fig. 5A-B, full western blot dataset in Supplemental Fig. S9). The prominent contribution of LRRC8C to VRAC-mediated glutamate release in mouse astrocytes was surprising and contrasted with previous reports in other cell types, including our own findings in rat astrocytes, which showed that LRRC8C knockdown does not meaningfully impact swelling-activated D-[^3^H]aspartate efflux (see (55) and Discussion).

**Figure 5:**
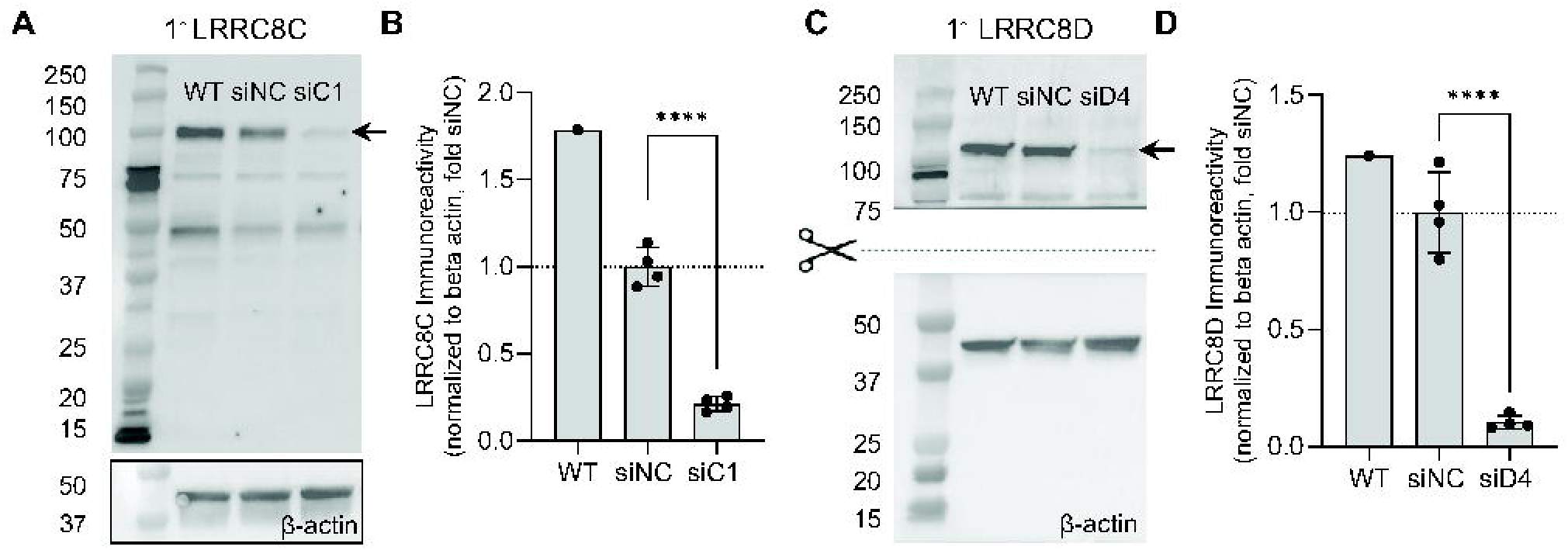
Validation of LRRC8C and LRRC8D protein loss following siRNA-mediated knockdown. **A**, Representative full-length Western blot of LRRC8C immunoreactivity and corresponding β-actin signal from protein lysates prepared 96h post siRNA transfection with either siNC or siC1. **B**, Quantification of LRRC8C protein levels normalized to β-actin and presented relative to the average signal from siNC samples. Data are the mean values ± SD from four independent transfections performed in four different astrocyte cultures. ****p < 0.0001, unpaired t-test. **C**, Representative Western blot of LRRC8D immunoreactivity and corresponding β-actin signal from protein lysates prepared 96h post siRNA transfection with either siNC or siD4. **D**, Quantification of LRRC8D protein levels normalized to β-actin and presented relative to the average signal from siNC samples. Data are the mean values ± SD from four independent transfections performed in four different astrocyte cultures. ***p < 0.001, unpaired t-test.

LRRC8D knockdowns produced more nuanced effects. Two of the tested siRNA constructs, siD2 and siD4, reduced VRAC activity by 10% and 28%, respectively (p = 0.542 and 0.002 vs. siNC, Fig. 4E2). In contrast, siD1 produced a striking 60% increase in swelling-activated D-[^3^H]aspartate release (p < 0.0001 vs. siNC; Fig. 4E2). Follow-up qRT-PCR analysis revealed that siD1 induced off-target effects, leading to a paradoxical upregulation of *Lrrc8a* and *Lrrc8c* transcript levels by 2.5-fold and 1.7-fold, respectively (Supplemental Fig. S10). Based on its superior efficacy and lack of off-target effects, siD4 was used in all subsequent experiments. Western blotting showed a ∼90% decrease in LRRC8D protein 96h post siRNA-transfection (p < 0.0001; Fig. 5C-D, full western blot dataset in Supplemental Fig. S11). Overall, these findings suggest that LRRC8D plays a role in glutamate-permeable VRAC function, though its contribution is relatively limited.

Finally, we tested three LRRC8E-targeting siRNA constructs (siE1, siE2, and siE4). Although the LRRC8E subunit is expressed in mouse astrocytes at only trace levels (see Fig. 2C), our prior work in rat astroglia has suggested that LRRC8E may be incorporated into astrocytic VRACs and modulate swelling-activated glutamate release (55). In the current study, LRRC8E knockdown had no biologically significant effect on VRAC activity in mouse astrocytes, albeit small increases in VRAC activity were observed with all siRNA constructs (p = 0.001 vs. siNC, Fig. 4F2).

### siRNA Knockdown of LRRC8 Isoforms Modifies Protein Levels of Partnering Subunits Without Affecting mRNA Expression

To assess siRNA specificity and determine whether loss of individual LRRC8 subunits influences the expression of the other family members, we performed qRT-PCR analysis of all five *Lrrc8* transcripts in astrocytes transfected with the most effective siRNA constructs targeting *Lrrc8a-d* (siA1, siB4, siC1, and siD4). At 48 hours post-transfection, knockdown of individual *Lrrc8* subunits significantly altered only the targeted transcripts, without a strong effect on the remaining *Lrrc8* genes (Fig. 6A-D). These results demonstrate the high specificity of this approach.

**Figure 6:**
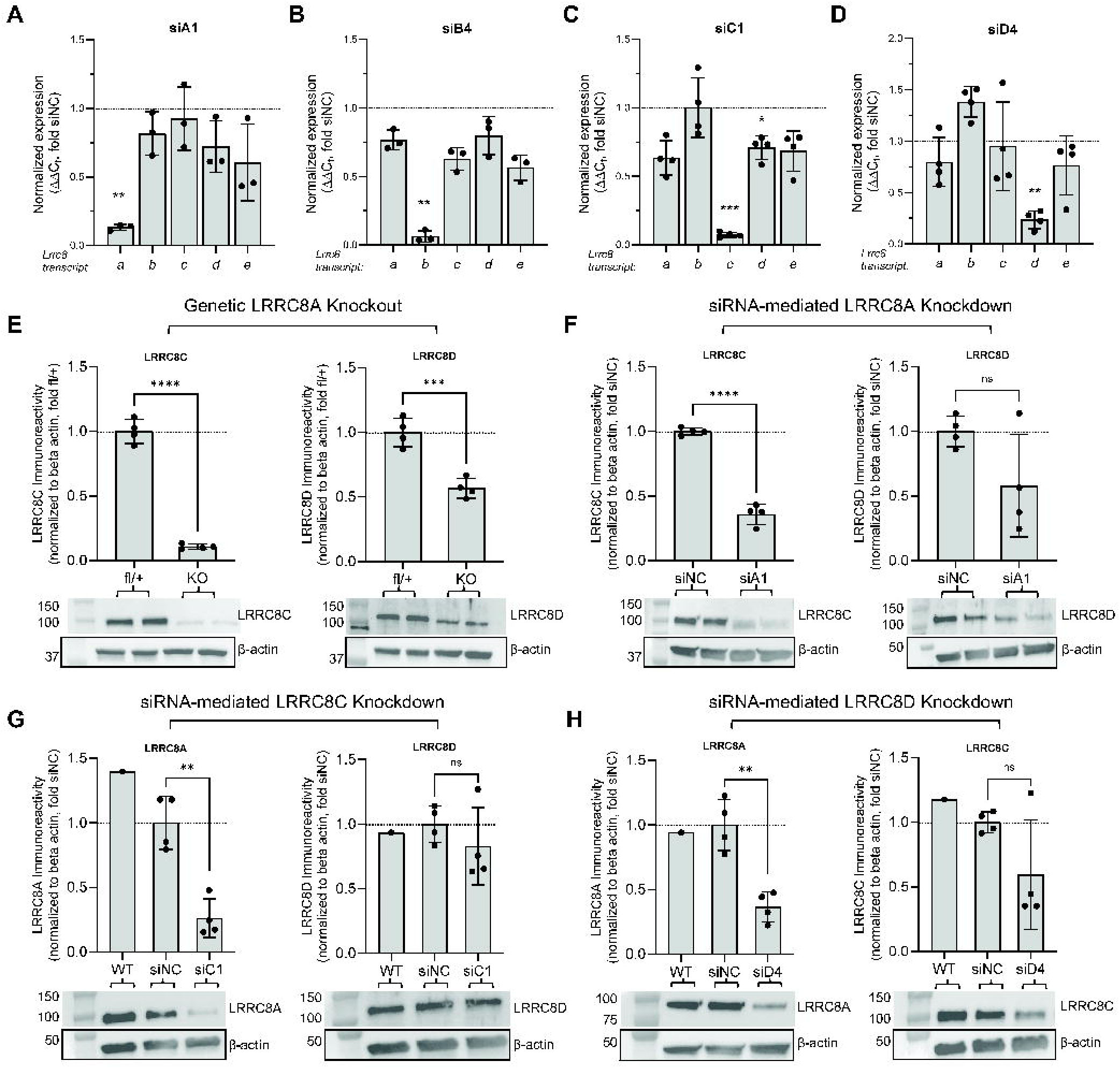
LRRC8 knockdown modulates partner protein levels independently of mRNA changes. **A-D**, qRT-PCR analysis of all five *Lrrc8* transcripts in mouse astrocytes treated with the most effective siRNA constructs targeting *Lrrc8a-d* (siA1, siB4, siC1, and siD4). Relative *Lrrc8a-e* expression levels are normalized to *Rpl13a* within the same sample and compared to siNC, measured 48 h post-transfection. Data are the mean normalized values ± SD of 3-4 independent transfections. ***p < 0.001, **p < 0.01, *p < 0.05, one-population t-test with Bonferroni correction for multiple comparisons. **E**, Top: Quantification of LRRC8C and LRRC8D protein levels in control (*Lrrc8a*^fl/+^) and LRRC8A KO astrocytes, normalized to β-actin and expressed relative to the mean signal in control samples. Data are the mean values ± SD from four independent cultures per group. ****p < 0.0001, ***p < 0.001; unpaired t-test. Bottom: Representative Western blot of LRRC8C or LRRC8D immunoreactivity and matching β-actin signal. **F**, Top: Quantification of LRRC8C and LRRC8D protein levels in astrocytes treated with siNC or siA1, normalized to β-actin and expressed relative to siNC. Data are the mean values ± SD from four independent cultures per group. ****p < 0.0001; unpaired t-test. Bottom: Representative Western blots of LRRC8C or LRRC8D and matching β-actin. **G**, Top: Quantification of LRRC8A and LRRC8D protein levels in astrocytes treated with siNC or siC1, normalized to β-actin and expressed relative to siNC. Data are the mean values ± SD from four independent cultures per group. **p < 0.01, unpaired t-test. Bottom: Representative Western blot of LRRC8A or LRRC8D immunoreactivity and matching β-actin signal. **H**, Top: Quantification of LRRC8A and LRRC8C protein levels in astrocytes treated with siNC or siD4, normalized to β-actin and expressed relative to siNC. Data are the mean values ± SD from four independent cultures per group. **p < 0.01; unpaired t-test. Bottom: Representative Western blot or LRRC8A or LRRC8C immunoreactivity and matching β-actin signal.

Given the established role of LRRC8A in membrane trafficking (24, 65), we next tested whether manipulation of LRRC8A affects the protein stability of the other isoforms. Western blot analysis of protein lysates from control (*Lrrc8a*^fl/+^) and LRRC8A KO cultures revealed that LRRC8A deletion significantly reduced LRRC8C and LRRC8D protein levels, with a more pronounced effect on LRRC8C (>80% reduction, p < 0.001; Fig. 6E) compared with LRRC8D (∼40% reduction, p < 0.001; Fig. 6E, full western blot dataset in Supplemental Fig. S12).

The LRRC8A KO effects on LRRC8C and LRRC8D protein levels were partially recapitulated in siA1-treated astrocytes. LRRC8A siRNA knockdown reduced LRRC8C protein levels by 64% (p < 0.0001, Fig. 6F) and LRRC8D levels by 42% (not significant, p = 0.089, Fig. 6F, full western blot dataset in Supplemental Fig. S13). Altogether, these data show that both constitutive deletion and siRNA-mediated knockdown of LRRC8A reduce the protein levels of the partnering LRRC8C and LRRC8D subunits without altering their mRNA levels.

To determine whether the regulation of protein stability is LRRC8A-specific, we tested the effects of siRNA-mediated knockdowns of LRRC8C and LRRC8D. LRRC8C depletion reduced LRRC8A immunoreactivity by ∼74% (p = 0.001, Fig. 6G, full western blot dataset in Supplemental Fig. S14). LRRC8D knockdown reduced LRRC8A protein levels by ∼63% (p = 0.002, Fig. 6H, full western blot dataset in Supplemental Fig. S15). In contrast, there was no strong mutual interdependence between LRRC8C and LRRC8D. Specifically, LRRC8C knockdown had no effect on LRRC8D (p = 0.34, Fig. 6G), and LRRC8D knockdown produced a ∼40% decrease in LRRC8C immunoreactivity, but this effect was not statistically significant (p = 0.11, Fig. 6H). The pattern of selective co-stability of LRRC8 subunits supports the existence of distinct LRRC8A/C-and LRRC8A/D-containing VRAC populations.

### Non-Redundant Functions of LRRC8C and LRRC8D in Glutamate-Permeable VRACs

Although LRRC8C knockdown potently suppressed swelling-activated D-[^3^H]aspartate release (Fig. 4D2), the resulting inhibition was less pronounced than that observed after knockdown of the essential LRRC8A (compare to Fig. 4B2). These findings may suggest that other LRRC8 proteins function redundantly with LRRC8C in forming heteromeric VRACs. Our early work established that, in rat astrocytes, LRRC8C and LRRC8D serve as mutually replaceable partners of LRRC8A in forming glutamate-permeable VRACs (55). To test if this redundancy extends to mouse astrocytes, we performed a double-knockdown of LRRC8C and LRRC8D.

We first verified that co-transfection with siRNAs targeting *Lrrc8c* (siC1) and *Lrrc8d* (siD4) effectively reduced target gene expression by ≥80%, both individually and in combination (Fig. 7A). Functional assays performed in a multi-well plate revealed that dual knockdown of LRRC8C and LRRC8D did not further decrease glutamate release as compared to LRRC8C knockdown alone (57% vs. 52% inhibition, p = 0.240, Fig. 7B). To assess VRAC activity with higher temporal resolution, we repeated the double-knockdown experiments using a Lucite superfusion system. In this format, LRRC8C knockdown significantly reduced the maximum rate of glutamate release by ∼40% (p < 0.0001, Fig. 7C), whereas LRRC8D knockdown produced a smaller, non-significant ∼20% decrease (p = 0.084, Fig. 7C). Combined knockdown of LRRC8C+D inhibited VRAC activity by ∼48% (p < 0.001, Fig. 7C); however, this effect was not significantly different from LRRC8C knockdown alone (p = 0.688; Fig. 7C). Consistent with this finding, analysis of the integral VRAC-mediated release under hypoosmotic conditions showed no significant difference between siC1 alone and the siC1+siD4 combination (p = 0.603, Fig. 7C). While differences in the degree of VRAC inhibition between plate-based and superfusion formats were observed, the lack of an additive effect between LRRC8C and LRRC8D knockdowns was consistent across both assays.

**Figure 7:**
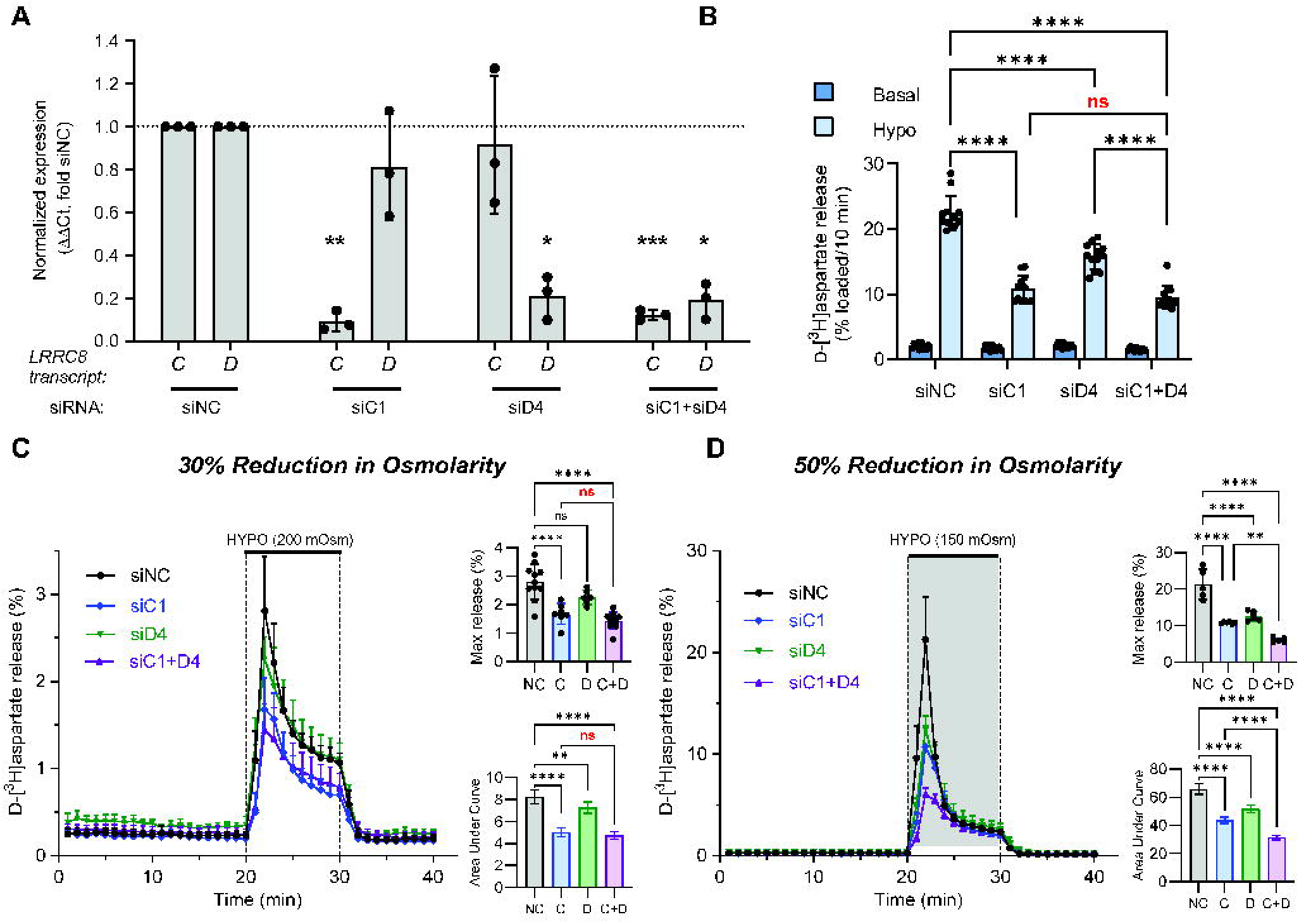
Non-redundant roles of LRRC8C and LRRC8D subunits in glutamate-permeable VRACs. **A**, Validation of siRNA efficacy in double knockdown experiments targeting *Lrrc8c* and *Lrrc8d*. Astrocytes were treated with siRNA targeting *Lrrc8c* (siC1), *Lrrc8d* (siD4), or their combination (siC1 + siD4). Efficacy of gene-specific knockdowns was verified by qRT-PCR. Data are the mean normalized expression values ± SD in three independently transfected cell cultures. **p < 0.01, *p < 0.05, one-population t-test with Bonferroni correction for multiple comparisons. **B**, Effect of siC1, siD4, or siC1+siD4 on VRAC-mediated D-[^3^H]aspartate release from cultured astrocytes exposed to a 30% reduction in medium osmolality measured in a 12-well plate format. Data are the mean values ± SD from twelve independent transfections in three different astrocyte cultures. ****p < 0.0001 vs. siNC, repeated measures two-way ANOVA with Tukey’s multiple comparisons test (only the effects of siRNAs under hypoosmotic conditions are shown). **C**, Kinetics of VRAC-mediated D-[^3^H]aspartate release from astrocyte cultures treated with siNC, siC1, siD4, or siC1+siD4 and exposed to a 30% reduction in medium osmolality. Data are the mean values ± SD in 6-11 independent transfections performed in three different astrocyte cultures. **Inset**: Statistical comparison of the maximal D-[^3^H]aspartate release values (top) and area under the curve (bottom) in the same experiments. ****p < 0.0001, ***p < 0.001, **p < 0.01, one-way ANOVA with Tukey’s multiple comparisons test. **D**, Kinetics of VRAC-mediated D-[^3^H]aspartate release from astrocytes treated with siNC, siC1, siD4, or siC1+siD4 and exposed to a 50% reduction in medium osmolality. Data are the mean values ± SD in 5-6 independent transfections performed in two different astrocyte cultures. **Inset**: Statistical comparison of the maximal D-[^3^H]aspartate release values (top) and area under the curve (bottom) in the same experiments. ****p < 0.0001, **p < 0.01, one-way ANOVA with Tukey’s multiple comparisons test.

Our LRRC8A KO data, particularly the heterozygous LRRC8A deletion results, suggest that in cells exposed to a 30% reduction in medium osmolality, VRAC channels may retain spare capacity, obscuring the full effects of LRRC8 subunit knockdown. Therefore, we repeated the double-knockdown experiments in astrocytes exposed to a 50% reduction in medium osmolarity (Fig. 7D). Under these conditions, knockdown of individual LRRC8C and LRRC8D subunits strongly inhibited VRAC activity (∼53% and ∼41% decrease in maximal release rates, respectively; p < 0.0001 vs. siNC, Fig. 7D). The double knockdown of LRRC8C plus LRRC8D produced partially additive ∼72% inhibition (p = 0.004, vs LRRC8C knockdown and p = 0.0001 vs. LRRC8D depletion alone, Fig. 7D). Partial additivity of the double-knockdown (as opposed to synergistic effect) is consistent with the existence of separate LRRC8A+LRRC8C and LRRC8A+ LRRC8D-containing heteromeric VRACs (see Discussion).

### LRRC8B Downregulation Rescues the Effects of LRRC8C- and LRRC8D-Knockdowns on VRAC Activity

Given that the combined downregulation of LRRC8C and LRRC8D did not reach the full effect of LRRC8A knockdown, we hypothesized that another abundant LRRC8 isoform, namely LRRC8B, serves as a functional substitute for either LRRC8C and/or LRRC8D. To test this hypothesis, we explored the potential contribution of LRRC8B in double-knockdown siRNA assays. Our prediction was that knocking down either LRRC8C+LRRC8B or LRRC8D+LRRC8B would produce a more substantial functional effect than knocking down LRRC8C or LRRC8D alone.

We first validated the specificity and efficacy of the double-knockdown of LRRC8B and LRRC8C. qRT-PCR analysis confirmed that siB4 and siC1 constructs were equally effective when transfected alone or in combination (Fig. 8A). Unexpectedly, in functional VRAC assays, the double knockdown of LRRC8B and LRRC8C resulted in a partial rescue of VRAC-mediated glutamate release as compared to the effect of LRRC8C knockdown on its own (Fig. 8B). Specifically, the siB4 construct had no effect (p = 0.98; Fig. 8B), siC1 suppressed VRAC activity by 51% (p < 0.0001; Fig. 8B), and the combination of siB4 and siC1 paradoxically caused a smaller 25% inhibition (p < 0.0001, siB1+C1 vs. siC1 alone, Fig. 8B). This functional rescue appears to be independent of protein stability given that LRRC8B knockdown caused a ∼27% reduction in LRRC8C protein (p = 0.044, Fig. 8C) and had no effect on LRRC8A (p = 0.641, Supplemental Fig. S16).

**Figure 8:**
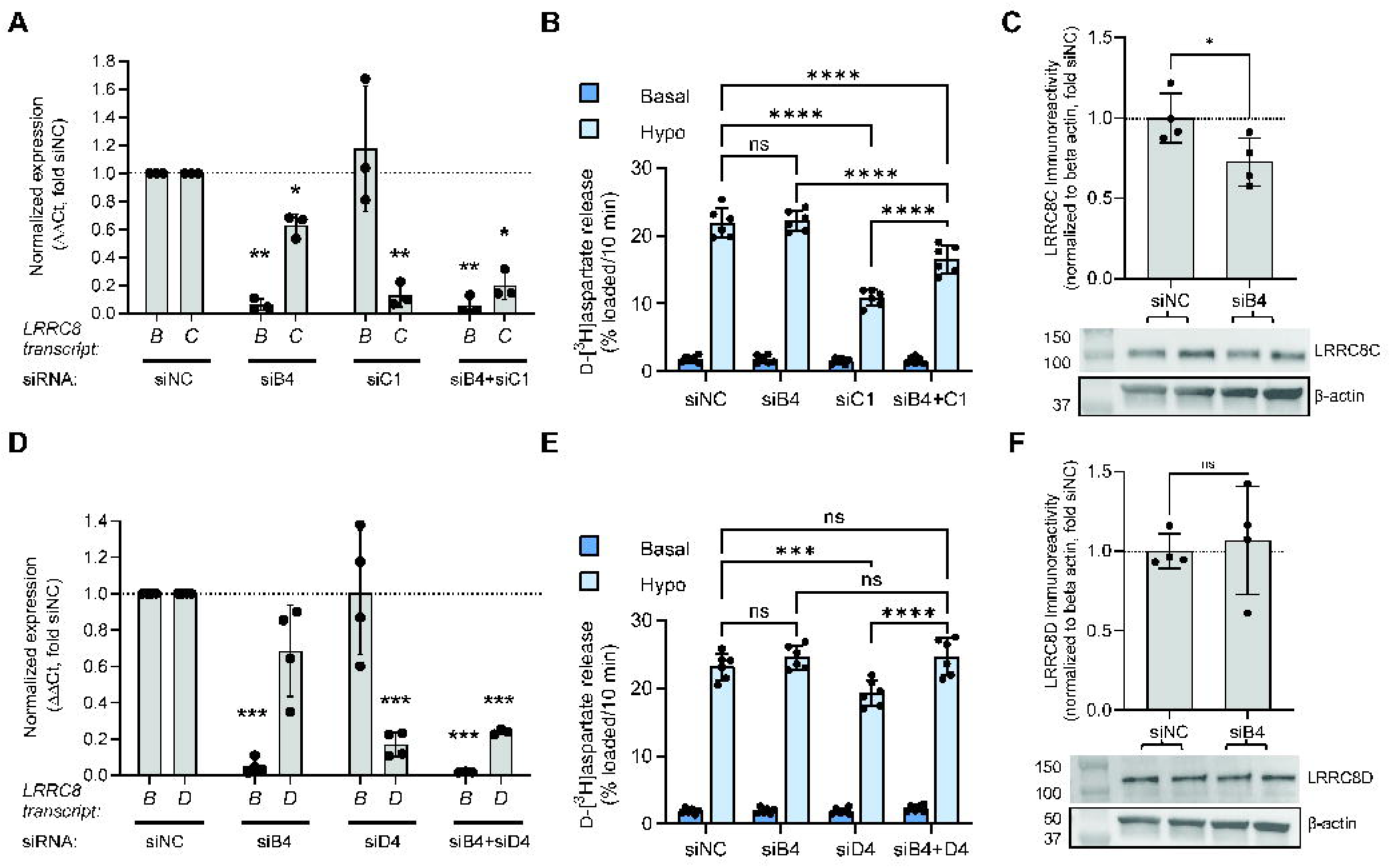
Functional contribution of LRRC8B to LRRC8C- and LRRC8D-containing VRACs. **A**, Validation of siRNA efficacy in double knockdown experiments targeting *Lrrc8b* and *Lrrc8c*. Astrocytes were treated with siRNA targeting *Lrrc8b* (siB4), *Lrrc8c* (siC1), or their combination (siB4+siC1). Efficacy of gene-specific knockdowns was verified by qRT-PCR. Data are the mean normalized expression values ± SD in three independently transfected cell cultures. **p < 0.01, *p < 0.05; one-population t-test with Bonferroni correction for multiple comparisons. **B**, Effects of siNC, siB4, siC1, siB4+siC1 on VRAC-mediated D-[^3^H]aspartate release from cultured astrocyte exposed to a 30% reduction in medium osmolality, measured in a 12-well plate format. Data are the mean values ± SD from six independent transfections in two different astrocyte cultures. ****p < 0.0001, repeated measures two-way ANOVA with Tukey’s multiple comparisons test (only the effects of siRNAs under hypoosmotic conditions are shown). **C**, Top: Quantification of LRRC8C protein levels in astrocytes treated with siNC or siB4, normalized to β-actin and expressed relative to siNC. Data are the mean values ± SD from four independent cultures per group. *p < 0.05; unpaired t-test. Bottom: Representative Western blot of LRRC8C and matching β-actin. **D**, qRT-PCR validation of double siRNA knockdown experiments targeting *Lrrc8b* (siB4) and *Lrrc8d* (siD4). Astrocytes were treated with siB4, siD4, or their combination (siB4+siD4). Data are the mean normalized expression values ± SD in three to four independently transfected cell cultures. ***p < 0.001, one-population t-test with Bonferroni correction for multiple comparisons. **E**, Effects of siNC, siB4, siD4, siB4+siD4 on VRAC-mediated D-[^3^H]aspartate release from cultured astrocyte exposed to a 30% reduction in medium osmolality measured in a 12-well plate format. Data are the mean values ± SD from six independent transfections in two different astrocyte cultures. ****p < 0.0001, ***p < 0.001; repeated measures two-way ANOVA with Tukey’s multiple comparisons test (only the effects of siRNAs under hypoosmotic conditions are shown). **F**, Top: Quantification of LRRC8D protein levels in astrocytes treated with siNC or siB4, normalized to β-actin and expressed relative to siNC. Data are the mean values ± SD from four independent cultures per group, unpaired t-test. Bottom: Representative Western blot LRRC8D and matching β-actin.

Similarly, we quantified the efficacy of double-knockdown of LRRC8B and LRRC8D. qRT-PCR analysis verified that siB4 and siD4 constructs were equally effective when transfected alone or in combination (Fig. 8D). In functional assays, the double knockdown of LRRC8B and LRRC8D resulted in a complete rescue of the VRAC-mediated glutamate release as compared to the effect LRRC8D knockdown on its own (Fig. 8E). Specifically, siB4 had no effect (p = 0.415; Fig. 8E), siD4 reduced VRAC activity by 17% (p < 0.001; Fig. 8E), and the combination of siB4 and siD4 caused full recovery of glutamate release in hypoosmotic medium (p = 0.36, Fig. 8E). In complementary Western blot analysis, we found no effects of LRRC8B knockdown on LRRC8D protein levels (p = 0.717, Fig. 8F, full western blot dataset in Supplemental Fig. S17).

It is important to note that the rescue effects of co-transfection with siB4 were subunit specific. In experiments involving the combined knockdown of LRRC8A+LRRC8B, the effect of siA1+siB4 was no different from the effect of siA1 alone (Supplemental Fig. S18A). Overall, these data suggest that LRRC8B may act as a structural partner within LRRC8A+LRRC8C and LRRC8A+LRRC8D-containing channels.

## DISCUSSION

The main objective of this study was to identify which LRRC8 isoforms contribute to the formation of glutamate-permeable VRACs in mouse astrocytes. Our key findings can be summarized as follows: (1) Genetic knockout of the essential LRRC8A subunit confirmed that LRRC8A-containing VRACs are the primary pathway for astrocytic swelling-activated glutamate release and account for ∼98% of the release under conditions of substantial cell swelling. (2) RNAi analysis supported a model in which glutamate-permeable VRACs strongly depend on LRRC8A and LRRC8C. Interestingly, LRRC8A and LRRC8C are reciprocally required for the protein stability of their partner subunit, without affecting their mRNA levels. (3) The LRRC8D isoform provided an additional contribution to swelling-activated glutamate release, but is likely present in a distinct subpopulation of VRACs. This model is supported by both functional glutamate release assays and the lack of mutual regulation of LRRC8C and LRRC8D protein stability.

Despite extensive work in the field, it remains debated whether VRACs are the primary pathway for swelling-activated glutamate release in astrocytes. Several alternative mechanisms have been proposed, including swelling-activated SLCO2A1-containing Maxi Cl⁻ channels (66, 67), the Tweety homolog proteins TTYH1-3 (68), and Ca^2+^-activated BEST1 channels (69, 70). Furthermore, astroglial Cx43 hemichannels (71) and the P2X_7_ receptor-activated pannexin complexes (72, 73) can also release amino acid gliotransmitters in response to various stimuli. Consistent with our prior work (30), genetic deletion of the essential VRAC subunit LRRC8A largely abolished swelling-activated glutamate release. A small, swelling-independent residual component persisted in LRRC8A-null cells, likely mediated by an alternative pathway(s) or cell lysis, but its quantitative contribution appeared minimal. LRRC8A KO astrocytes also showed reduced basal glutamate release under isoosmotic conditions, indicating tonic VRAC activity in non-swollen cells. Interestingly, heterozygous LRRC8A deletion produced only a mild inhibition of glutamate release in astrocytes exposed to a 30% reduction in medium osmolarity [this study and (30)], but strongly suppressed glutamate efflux when cells were subjected to more substantial, supraphysiological cell swelling. One interpretation of these findings is that severe swelling activates the full complement of VRACs in the plasma membrane and recruits additional channels from intracellular organellar reserves, unmasking the maximal functional deficit. Together, these findings indicate that LRRC8A-containing VRACs dominate swelling-activated glutamate release and contribute to tonic glutamate efflux in non-swollen astrocytes.

The subunit composition of VRACs that conduct organic anions, particularly glutamate and aspartate, remains uncertain. Model biophysical studies have produced somewhat conflicting results. In CRISPR/Cas9-edited LRRC8-null HEK293 cells, heterologous co-expression of LRRC8A with LRRC8D or LRRC8A with LRRC8E produced glutamate-permeable VRACs, whereas complexes of LRRC8A+LRRC8C showed limited glutamate conductance (54). In comparison, in *Xenopus* oocytes, co-expression of LRRC8A with LRRC8C or LRRC8A with LRRC8E resulted in significant glutamate permeability, whereas LRRC8A+LRRC8D heteromers yielded smaller glutamate conductance (56). In native environments, VRACs are thought to contain multiple, functionally redundant LRRC8 subunits, since deletion of individual LRRC8B-E proteins failed to reduce endogenous VRAC currents in HEK293 or HeLa cells (24, 74). Consistent with this idea, we previously reported that, in rat astrocytes, endogenous glutamate-permeable VRACs are formed with redundant contributions from LRRC8C, LRRC8D, and LRRC8E (55). Within this context, we were surprised to find that in mouse astroglia, swelling-activated glutamate release strongly depended on LRRC8C expression, with 56% inhibition of glutamate release following individual LRRC8C knockdown. In contrast, downregulation of LRRC8D produced only a mild reduction in swelling-activated glutamate release, suggesting a more limited role for this isoform. Recent studies further highlight the importance of LRRC8C, demonstrating loss of VRAC currents in LRRC8C-null mouse T lymphocytes and marked inhibition of VRAC currents after LRRC8C knockdown in human umbilical endothelial cells (75, 76).

Given that depletion of either LRRC8C or LRRC8D alone failed to reproduce the strong inhibition observed after loss of LRRC8A, we next examined whether combined LRRC8C+ LRRC8D knockdown produced additive or synergistic inhibition. A synergistic effect, as previously reported in rat astrocytes (55), would support redundant contributions of LRRC8C and LRRC8D within a single heteromeric VRAC. Alternatively, additive effects would favor the existence of two distinct channel populations. In moderately swollen cells, combined knockdown of LRRC8C and LRRC8D did not further inhibit glutamate release beyond that observed with LRRC8C depletion alone (Fig. 9). The clear absence of synergism argues against a single-channel model. Although the lack of additivity introduces some uncertainty, the glutamate release data are generally consistent with a two-channel model. The idea of two independent channels is further supported by the analysis of subunit-specific regulation of LRRC8 protein stability. We found strong reciprocal regulation between LRRC8A and LRRC8C, as well as between LRRC8A and LRRC8D. In contrast, we found minimal cross-regulation between LRRC8C and LRRC8D. The surprising interdependence of LRRC8 subunit stability has a recent precedent in the mouse endothelium, where LRRC8A, LRRC8B, and LRRC8C all mutually regulate the protein abundance of their partnering isoforms (76). We speculate that loss of a given VRAC subunit promotes degradation of its partners within the same channel complex, and the lack of co-dependence between LRRC8C and LRRC8D points to their presence in distinct VRAC assemblies.

**Figure 9:**
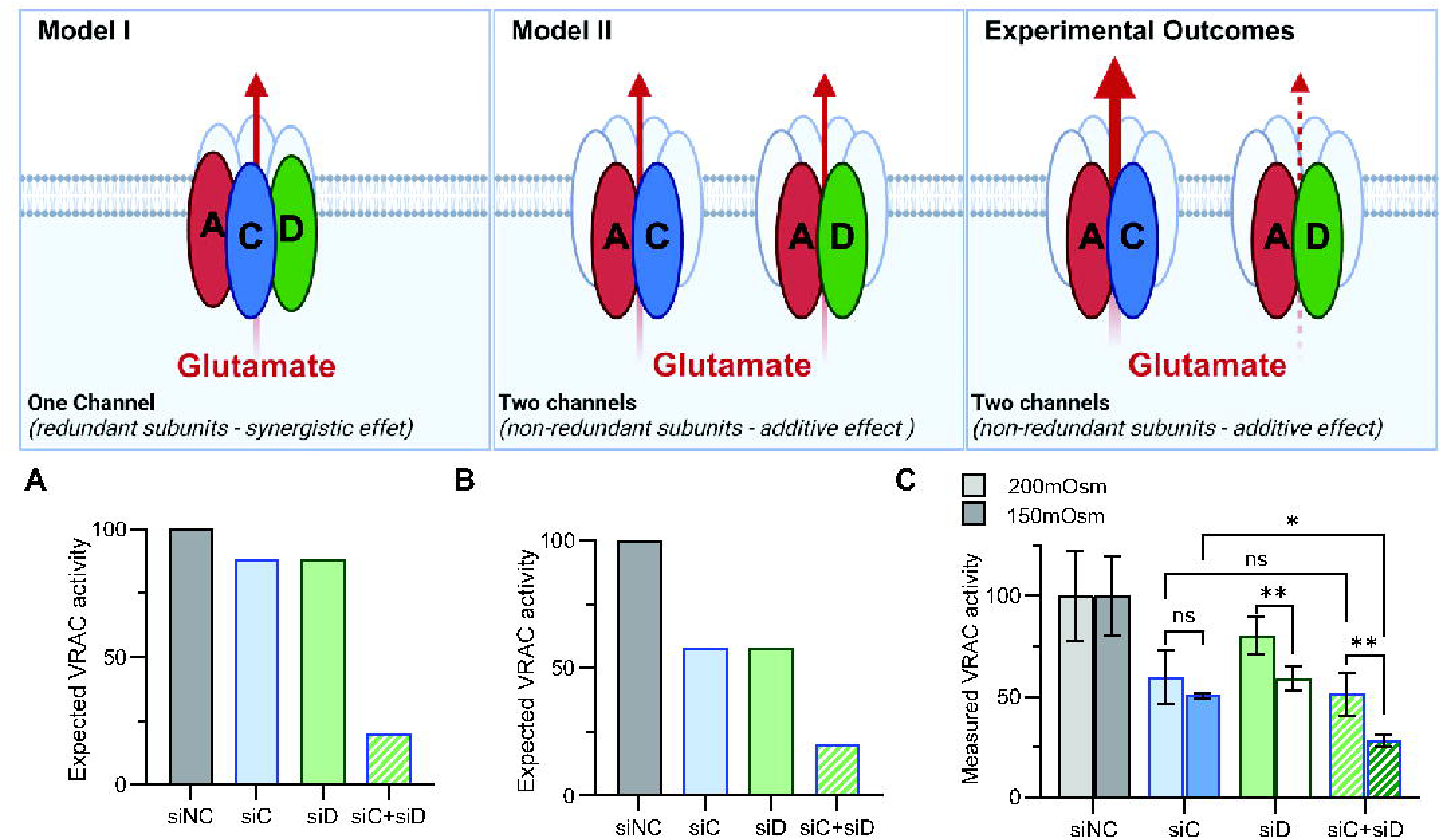
Proposed subunit composition of glutamate-permeable VRACs in astrocytes. **A** and **B**, Theoretical predictions of the effects on siRNA knockdowns on VRAC activity based on two models of VRAC composition. In *Model I*, VRACs are assembled from LRRC8A, LRRC8C, and LRRC8D subunits, with LRRC8C and LRRC8D functioning as mutually redundant components. In *Model II*, two independent VRAC populations are formed by either LRRC8A+LRRC8C or LRRC8A+LRRC8D. In this case, LRRC8C and LRRC8D subunits are not functionally redundant. **C**, *Experimental outcomes* showing the effects of LRRC8C (siC) and/or LRRC8D (siD) knockdown on VRAC-mediated D-[^3^H]aspartate release in mouse astrocytes subjected to moderate (30% reduction in osmolarity; light bars) or severe (50% reduction in osmolarity; dark bars) hypotonic stress (data from Fig. 7C and 7D). These results are consistent with the Model II in which two distinct VRAC populations mediate glutamate permeability. LRRC8A+LRRC8D-containing VRACs provide a smaller contribution to swelling-activated glutamate release, and their activity is dependent on the degree of cellular swelling. **p < 0.01, *p < 0.05; two-way ANOVA with Tukey’s multiple comparisons test.

Interestingly, the relative impact of LRRC8D depletion on swelling-activated glutamate release is substantially increased in astrocytes subjected to supraphysiological cell swelling, induced by a 50% reduction in medium osmolarity. In this instance, the combined knockdown of LRRC8C and LRRC8D produced nearly additive inhibition, consistent with a two-population model. The increased role of LRRC8D during severe cell swelling may reflect recruitment of additional LRRC8D-containing channels to the plasma membrane. In support of this idea, Groulx et al. reported that mammalian cells accommodated moderate cell volume increases primarily through shape changes, whereas in substantially swollen cells, membrane tension was compensated for by endomembrane insertion (77). While the organellar distribution of VRACs is incompletely understood, a substantial fraction of LRRC8 channels are thought to reside in the lysosomal compartment and are recruited to the plasmalemma in response to severe hypoosmotic stress (78). Also of importance, a recent study demonstrated that native lysosomal VRACs are composed of LRRC8A, LRRC8B, LRRC8D, and LRRC8E subunits, with a notable absence of LRRC8C (79). Thus, mobilization from intracellular pools is likely to preferentially enrich the surface expression of LRRC8D-containing VRACs.

Finally, we would like to address the functional role of LRRC8B, which is highly expressed in the mouse brain [(25) and present data]. In mouse astrocytes, LRRC8B knockdown alone did not inhibit swelling-activated glutamate release. However, the combined downregulation of LRRC8B in conjunction with LRRC8C or LRRC8D partially rescued the inhibitory effects of LRRC8C or LRRC8D depletion alone. Similar paradoxical observations have been reported in HEK293 cells, where deletion of LRRC8B from native VRACs increased swelling-activated D-[^3^H]aspartate efflux (54). Although the LRRC8B subunit can be incorporated into VRACs (24, 76), its relevance to VRAC activity remains unclear. To date, the only established role for LRRC8B is the modulation of Ca^2+^ leak from the endoplasmic reticulum (80). Heterologous co-expression of LRRC8A and LRRC8B fails to produce functional VRAC currents (24, 54, 56, 74). It is known that homomeric LRRC8A channels have little to no conductance due to steric restrictions, including the ring of positively charged arginine residues (R103) within the narrowest region of the pore (47-49). LRRC8B shares structural homology with LRRC8A and contains a matching arginine residue (R99), whereas LRRC8C-E have neutral amino acids at this position (47, 48, 50, 81). We speculate that substituting LRRC8B for LRRC8C or LRRC8D within VRAC heteromers partially narrows the channel pore, and that, in the double knockdown experiments, removing LRRC8B makes the residual VRACs more permeable to glutamate. It is also possible that LRRC8B knockdown effects may stem from modified intracellular Ca^2+^ signaling.

In summary, our findings establish that glutamate permeability of native VRACs in mouse astrocytes is largely determined by the non-redundant contributions of two subunits: the essential LRRC8A and the complementary LRRC8C. This refined understanding of VRAC composition in the brain has broad clinical relevance, given that both loss and gain of VRAC function produce severe neurological phenotypes in humans. For example, gain-of-function mutations in *LRRC8C* cause a severe multisystem disorder, termed *LRRC8C*-related TIMES syndrome, which is associated with microcephaly, intellectual disability, and seizures (82). Conversely, mutations causing megalencephalic leukoencephalopathy with subcortical cysts (MLC) indirectly reduce VRAC activity and cause macrocephaly, white-matter disorganization, and severe seizures (83-85). These neurological manifestations may, at least in part, be due to altered neurotransmitter release via VRACs. Defining the structural determinants that govern VRAC neurotransmitter permeability, therefore, provides critical insight into brain (patho)physiology and may guide the development of subunit-targeted VRAC therapies.

## DATA AVAILABILITY

All primary data included in this manuscript have been archived in the Open Science Framework depository (osf.io) and can be accessed at https://osf.io/xvyba. All original Western blot images are available in FigShare at https://doi.org/10.6084/m9.figshare.31143022

## SUPPLEMENTAL MATERIAL

Supplemental Figs. S1-S18: https://doi.org/10.6084/m9.figshare.31143022

## ACKNOWLEDGMENTS

The authors thank Dr. Preeti Dohare for help with validating LRRC8D siRNA constructs, Dr. Alena Rudkouskya for help with LRRC8A Western blot experiments, and Antonio M. Fidaleo for his advice on performing RNA-seq analyses. Illustrations in Figs. 2, 4, 9, and Graphical Abstract have been created with BioRender.

## GRANTS

This study was supported by the National Institutes of Health, grants R01 NS111943 and R21 NS140782 (to A.A.M.).

## DISCLOSURES

The authors have no conflict of interest, financial or otherwise, to disclose.

## AUTHOR CONTRIBUTIONS

Conceived and designed research: M.L.C. and A.A.M. Performed experiments: M.L.C., A.D.S., J.W.N. Analyzed data: M.L.C., A.D.S., A.A.M. Interpreted results: M.L.C. and A.A.M. Prepared figures: M.L.C. and A.A.M. Drafted manuscript: M.L.C. and A.A.M. Edited and revised manuscript, M.L.C., A.D.S., J.W.N., and A.A.M. Approved final version of manuscript: M.L.C., A.D.S., J.W.N., and A.A.M.

## Notes

https://osf.io/xvyba

https://doi.org/10.6084/m9.figshare.31143022

### Competing Interest Statement

The authors have declared no competing interest.

### Summary of Updates

This updated version of the manuscript contains three additional figures and textual revisions that address concerns raised in the peer-review process.

## REFERENCES

1. Strange K, Emma F and Jackson PS. Cellular and molecular physiology of volume-sensitive anion channels. Am J Physiol 270: C711–C730, 1996.

2. Okada Y. Volume expansion-sensing outward-rectifier Cl- channel: fresh start to the molecular identity and volume sensor. Am J Physiol 273: C755–C789, 1997. doi: 10.1152/ajpcell.1997.273.3.C755

3. Nilius B, Eggermont J, Voets T, Buyse G, Manolopoulos V and Droogmans G. Properties of volume-regulated anion channels in mammalian cells. Prog Biophys Mol Biol 68: 69–119, 1997.

4. Hazama A and Okada Y. Ca2+ sensitivity of volume-regulatory K+ and Cl- channels in cultured human epithelial cells. J Physiol 402: 687–702, 1988.

5. Cahalan MD and Lewis RS. Role of potassium and chloride channels in volume regulation by T lymphocytes. Soc Gen Physiol Ser 43: 281–301, 1988. doi: n/a

6. Worrell RT, Butt AG, Cliff WH and Frizzell RA. A volume-sensitive chloride conductance in human colonic cell line T84. Am J Physiol 256: C1111–C1119, 1989.

7. Solc CK and Wine JJ. Swelling-induced and depolarization-induced Cl- channels in normal and cystic fibrosis epithelial cells. Am J Physiol 261: C658–C674, 1991. doi: 10.1152/ajpcell.1991.261.4.C658

8. Kimelberg HK, Goderie SK, Higman S, Pang S and Waniewski RA. Swelling-induced release of glutamate, aspartate, and taurine from astrocyte cultures. J Neurosci 10: 1583–1591, 1990.

9. Roy G and Malo C. Activation of amino acid diffusion by a volume increase in cultured kidney (MDCK) cells. J Membr Biol 130: 83–90, 1992.

10. Kirk K, Ellory JC and Young JD. Transport of organic substrates via a volume-activated channel. J Biol Chem 267: 23475–23478, 1992.

11. Jackson PS and Strange K. Volume-sensitive anion channels mediate swelling-activated inositol and taurine efflux. Am J Physiol 265: C1489–C1500, 1993. doi: 10.1152/ajpcell.1993.265.6.C1489

12. Lang F, Busch GL, Ritter M, Volkl H, Waldegger S, Gulbins E and Haussinger D. Functional significance of cell volume regulatory mechanisms. Physiol Rev 78: 247–306, 1998.

13. Hoffmann EK, Lambert IH and Pedersen SF. Physiology of cell volume regulation in vertebrates. Physiol Rev 89: 193–277, 2009.

14. Eggermont J, Trouet D, Carton I and Nilius B. Cellular function and control of volume-regulated anion channels. Cell Biochem Biophys 35: 263–274, 2001.

15. Mongin AA. Volume-regulated anion channel--a frenemy within the brain. Pflugers Arch 468: 421–441, 2016.

16. Jentsch TJ. VRACs and other ion channels and transporters in the regulation of cell volume and beyond. Nat Rev Mol Cell Biol 17: 293–307, 2016.

17. Chen L, Konig B, Liu T, Pervaiz S, Razzaque YS and Stauber T. More than just a pressure relief valve: physiological roles of volume-regulated LRRC8 anion channels. Biol Chem 400: 1481–1496, 2019. doi: 10.1515/hsz-2019-0189

18. Okada Y. Physiology of the volume-sensitive/regulatory anion channel VSOR/VRAC: part 2: its activation mechanisms and essential roles in organic signal release. J Physiol Sci 74: 34, 2024. doi: 10.1186/s12576-023-00897-x

19. Thone FMB, Polovitskaya MM and Jentsch TJ. LRRC8/VRAC chloride and metabolite channels in signaling and volume regulation. Trends Biochem Sci 50: 873–891, 2025. doi: 10.1016/j.tibs.2025.07.001

20. Pedersen SF, Okada Y and Nilius B. Biophysics and physiology of the volume-regulated anion channel (VRAC)/volume-sensitive outwardly rectifying anion channel (VSOR). Pflugers Arch 468: 371–383, 2016.

21. Strange K, Yamada T and Denton JS. A 30-year journey from volume-regulated anion currents to molecular structure of the LRRC8 channel. J Gen Physiol 151: 100–117, 2019.

22. Okada Y. Physiology of the volume-sensitive/regulatory anion channel VSOR/VRAC. Part 1: from its discovery and phenotype characterization to the molecular entity identification. J Physiol Sci 74: 3, 2024. doi: 10.1186/s12576-023-00897-x

23. Qiu Z, Dubin AE, Mathur J, Tu B, Reddy K, Miraglia LJ, Reinhardt J, Orth AP and Patapoutian A. SWELL1, a plasma membrane protein, is an essential component of volume-regulated anion channel. Cell 157: 447–458, 2014.

24. Voss FK, Ullrich F, Munch J, Lazarow K, Lutter D, Mah N, Andrade-Navarro MA, von Kries JP, Stauber T and Jentsch TJ. Identification of LRRC8 heteromers as an essential component of the volume-regulated anion channel VRAC. Science 344: 634–638, 2014. doi: 10.1126/science.1252826

25. Abascal F and Zardoya R. LRRC8 proteins share a common ancestor with pannexins, and may form hexameric channels involved in cell-cell communication. Bioessays 34: 551–560, 2012.

26. Kumar L, Chou J, Yee CS, Borzutzky A, Vollmann EH, von Andrian UH, Park SY, Hollander G, Manis JP, Poliani PL and Geha RS. Leucine-rich repeat containing 8A (LRRC8A) is essential for T lymphocyte development and function. J Exp Med 211: 929–942, 2014.

27. Zhang Y, Xie L, Gunasekar SK, Tong D, Mishra A, Gibson WJ, Wang C, Fidler T, Marthaler B, Klingelhutz A, Abel ED, Samuel I, Smith JK, Cao L and Sah R. SWELL1 is a regulator of adipocyte size, insulin signalling and glucose homeostasis. Nat Cell Biol 19: 504–517, 2017.

28. Yang J, Vitery MDC, Chen J, Osei-Owusu J, Chu J and Qiu Z. Glutamate-releasing SWELL1 channel in astrocytes modulates synaptic transmission and promotes brain damage in stroke. Neuron 102: 813–827, 2019. doi: 10.1016/j.neuron.2019.03.029

29. Kumar A, Xie L, Ta CM, Hinton AO, Gunasekar SK, Minerath RA, Shen K, Maurer JM, Grueter CE, Abel ED, Meyer G and Sah R. SWELL1 regulates skeletal muscle cell size, intracellular signaling, adiposity and glucose metabolism. eLife 9: 2020.

30. Wilson CS, Dohare P, Orbeta S, Nalwalk JW, Huang Y, Ferland RJ, Sah R, Scimemi A and Mongin AA. Late adolescence mortality in mice with brain-specific deletion of the volume-regulated anion channel subunit LRRC8A. FASEB J 35: e21869, 2021. doi: 10.1096/fj.202002745R

31. Zhou JJ, Luo Y, Chen SR, Shao JY, Sah R and Pan HL. LRRC8A-dependent volume-regulated anion channels contribute to ischemia-induced brain injury and glutamatergic input to hippocampal neurons. Exp Neurol 332: 113391, 2020.

32. Kimelberg HK, MacVicar BA and Sontheimer H. Anion channels in astrocytes: Biophysics, pharmacology, and function. Glia 54: 747–757, 2006.

33. Akita T and Okada Y. Characteristics and roles of the volume-sensitive outwardly rectifying (VSOR) anion channel in the central nervous system. Neuroscience 275: 211–231, 2014.

34. Wilson CS and Mongin AA. Cell volume control in healthy brain and neuropathologies. Curr Top Membr 81: 385–455, 2018.

35. Kimelberg HK. Astrocytic swelling in cerebral ischemia as a possible cause of injury and target for therapy. Glia 50: 389–397, 2005.

36. Mongin AA. Disruption of ionic and cell volume homeostasis in cerebral ischemia: The perfect storm. Pathophysiology 14: 183–193, 2007.

37. Mongin AA and Kimelberg HK. ATP potently modulates anion channel-mediated excitatory amino acid release from cultured astrocytes. Am J Physiol Cell Physiol 283: C569–C578, 2002.

38. Takano T, Kang J, Jaiswal JK, Simon SM, Lin JH, Yu Y, Li Y, Yang J, Dienel G, Zielke HR and Nedergaard M. Receptor-mediated glutamate release from volume sensitive channels in astrocytes. Proc Natl Acad Sci U S A 102: 16466–16471, 2005.

39. Liu HT, Akita T, Shimizu T, Sabirov RZ and Okada Y. Bradykinin-induced astrocyte-neuron signalling: glutamate release is mediated by ROS-activated volume-sensitive outwardly rectifying anion channels. J Physiol 587: 2197–2209, 2009.

40. Yang J, Chen J, Liu Y, Chen KH, Baraban JM and Qiu Z. Ventral tegmental area astrocytes modulate cocaine reward by tonically releasing GABA. Neuron 111: 1104–1117, 2023. doi: 10.1016/j.neuron.2022.12.033

41. Kimelberg HK, Feustel PJ, Jin Y, Paquette J, Boulos A, Keller RW, Jr. and Tranmer BI. Acute treatment with tamoxifen reduces ischemic damage following middle cerebral artery occlusion. Neuroreport 11: 2675–2679, 2000.

42. Kimelberg HK, Jin Y, Charniga C and Feustel PJ. Neuroprotective activity of tamoxifen in permanent focal ischemia. J Neurosurg 99: 138–142, 2003.

43. Zhang Y, Zhang H, Feustel PJ and Kimelberg HK. DCPIB, a specific inhibitor of volume regulated anion channels (VRACs), reduces infarct size in MCAo and the release of glutamate in the ischemic cortical penumbra. Exp Neurol 210: 514–520, 2008.

44. Boulos AS, Deshaies EM, Dalfino JC, Feustel PJ, Popp AJ and Drazin D. Tamoxifen as an effective neuroprotectant in an endovascular canine model of stroke. J Neurosurg 114: 1117–1126, 2011.

45. Chen J, Yang J, Chu J, Chen KH, Alt J, Rais R and Qiu Z. The SWELL1 channel promotes ischemic brain damage by mediating neuronal swelling and glutamate toxicity. Adv Sci (Weinh) 11: e2401085, 2024. doi: 10.1002/advs.202401085

46. Balkaya M, Dohare P, Chen S, Schober AL, Fidaleo AM, Nalwalk JW, Sah R and Mongin AA. Conditional deletion of LRRC8A in the brain reduces stroke damage independently of swelling-activated glutamate release. iScience 26: 106669, 2023. doi: 10.1016/j.isci.2023.106669

47. Deneka D, Sawicka M, Lam AKM, Paulino C and Dutzler R. Structure of a volume-regulated anion channel of the LRRC8 family. Nature 558: 254–259, 2018.

48. Kasuya G, Nakane T, Yokoyama T, Jia Y, Inoue M, Watanabe K, Nakamura R, Nishizawa T, Kusakizako T, Tsutsumi A, Yanagisawa H, Dohmae N, Hattori M, Ichijo H, Yan Z, Kikkawa M, Shirouzu M, Ishitani R and Nureki O. Cryo-EM structures of the human volume-regulated anion channel LRRC8. Nat Struct Mol Biol 25: 797–804, 2018.

49. Kefauver JM, Saotome K, Dubin AE, Pallesen J, Cottrell CA, Cahalan SM, Qiu Z, Hong G, Crowley CS, Whitwam T, Lee WH, Ward AB and Patapoutian A. Structure of the human volume regulated anion channel. eLife 7: e38461, 2018.

50. Kern DM, Oh S, Hite RK and Brohawn SG. Cryo-EM structures of the DCPIB-inhibited volume-regulated anion channel LRRC8A in lipid nanodiscs. eLife 8: e42636, 2019. doi: 10.7554/eLife.42636

51. Rutz S, Deneka D, Dittmann A, Sawicka M and Dutzler R. Structure of a volume-regulated heteromeric LRRC8A/C channel. Nat Struct Mol Biol 30: 52–61, 2023.

52. Lurie A, Stephens CA, Kern DM, Henn KM, Latorraca NR and Brohawn SG. Assembly and lipid-gating of LRRC8A:D volume-regulated anion channels. Nat Commun 17: 366, 2026. doi: 10.1038/s41467-025-67052-5

53. Planells-Cases R, Lutter D, Guyader C, Gerhards NM, Ullrich F, Elger DA, Kucukosmanoglu A, Xu G, Voss FK, Reincke SM, Stauber T, Blomen VA, Vis DJ, Wessels LF, Brummelkamp TR, Borst P, Rottenberg S and Jentsch TJ. Subunit composition of VRAC channels determines substrate specificity and cellular resistance to Pt-based anti-cancer drugs. EMBO J 34: 2993–3008, 2015. doi: 10.15252/embj.201592409

54. Lutter D, Ullrich F, Lueck JC, Kempa S and Jentsch TJ. Selective transport of neurotransmitters and modulators by distinct volume-regulated LRRC8 anion channels. J Cell Sci 130: 1122–1133, 2017. doi: 10.1242/jcs.196253

55. Schober AL, Wilson CS and Mongin AA. Molecular composition and heterogeneity of the LRRC8-containing swelling-activated osmolyte channels in primary rat astrocytes. J Physiol 595: 6939–6951, 2017. doi: 10.1113/JP275053

56. Gaitan-Penas H, Gradogna A, Laparra-Cuervo L, Solsona C, Fernandez-Duenas V, Barrallo-Gimeno A, Ciruela F, Lakadamyali M, Pusch M and Estevez R. Investigation of LRRC8-mediated volume-regulated anion currents in Xenopus oocytes. Biophys J 111: 1429–1443, 2016. doi: 10.1016/j.bpj.2016.08.030

57. Livak KJ and Schmittgen TD. Analysis of relative gene expression data using real-time quantitative PCR and the 2(-Delta Delta C(T)) Method. Methods 25: 402–408, 2001.

58. Schneider CA, Rasband WS and Eliceiri KW. NIH Image to ImageJ: 25 years of image analysis. Nat Methods 9: 671–675, 2012.

59. Abdullaev IF, Rudkouskaya A, Schools GP, Kimelberg HK and Mongin AA. Pharmacological comparison of swelling-activated excitatory amino acid release and Cl- currents in rat cultured astrocytes. J Physiol 572: 677–689, 2006. doi: 10.1113/jphysiol.2005.103820

60. Bowens NH, Dohare P, Kuo YH and Mongin AA. DCPIB, the proposed selective blocker of volume-regulated anion channels, inhibits several glutamate transport pathways in glial cells. Mol Pharmacol 83: 22–32, 2013. doi: 10.1124/mol.112.080457

61. Schober AL and Mongin AA. Intracellular levels of glutamate in swollen astrocytes are preserved via neurotransmitter reuptake and de novo synthesis: implications for hyponatremia. J Neurochem 135: 176–185, 2015. doi: 10.1111/jnc.13229

62. Tronche F, Kellendonk C, Kretz O, Gass P, Anlag K, Orban PC, Bock R, Klein R and Schutz G. Disruption of the glucocorticoid receptor gene in the nervous system results in reduced anxiety. Nat Genet 23: 99–103, 1999. doi: 10.1038/12703

63. Dubois NC, Hofmann D, Kaloulis K, Bishop JM and Trumpp A. Nestin-Cre transgenic mouse line Nes-Cre1 mediates highly efficient Cre/loxP mediated recombination in the nervous system, kidney, and somite-derived tissues. Genesis 44: 355–360, 2006.

64. Verkhratsky A, Krishtal OA and Burnstock G. Purinoceptors on neuroglia. Mol Neurobiol 39: 190–208, 2009.

65. Yamada T and Strange K. Intracellular and extracellular loops of LRRC8 are essential for volume-regulated anion channel function. J Gen Physiol 150: 1003–1015, 2018. doi: 10.1085/jgp.201812016

66. Liu HT, Tashmukhamedov BA, Inoue H, Okada Y and Sabirov RZ. Roles of two types of anion channels in glutamate release from mouse astrocytes under ischemic or osmotic stress. Glia 54: 343–357, 2006. doi: 10.1002/glia.20400

67. Sabirov RZ, Merzlyak PG, Okada T, Islam MR, Uramoto H, Mori T, Makino Y, Matsuura H, Xie Y and Okada Y. The organic anion transporter SLCO2A1 constitutes the core component of the Maxi-Cl channel. EMBO J 36: 3309–3324, 2017.

68. Han YE, Kwon J, Won J, An H, Jang MW, Woo J, Lee JS, Park MG, Yoon BE, Lee SE, Hwang EM, Jung JY, Park H, Oh SJ and Lee CJ. Tweety-homolog (Ttyh) family encodes the pore-forming subunits of the swelling-dependent volume-regulated anion channel (VRAC(swell)) in the brain. Exp Neurobiol 28: 183–215, 2019. doi: 10.5607/en.2019.28.2.183

69. Woo DH, Han KS, Shim JW, Yoon BE, Kim E, Bae JY, Oh SJ, Hwang EM, Marmorstein AD, Bae YC, Park JY and Lee CJ. TREK-1 and Best1 channels mediate fast and slow glutamate release in astrocytes upon GPCR activation. Cell 151: 25–40, 2012.

70. Oh SJ, Han KS, Park H, Woo DH, Kim HY, Traynelis SF and Lee CJ. Protease activated receptor 1-induced glutamate release in cultured astrocytes is mediated by Bestrophin-1 channel but not by vesicular exocytosis. Mol Brain 5: 38, 2012.

71. Ye ZC, Wyeth MS, Baltan-Tekkok S and Ransom BR. Functional hemichannels in astrocytes: a novel mechanism of glutamate release. J Neurosci 23: 3588–3596, 2003.

72. Duan S, Anderson CM, Keung EC, Chen Y, Chen Y and Swanson RA. P2X7 receptor-mediated release of excitatory amino acids from astrocytes. J Neurosci 23: 1320–1328, 2003.

73. Iglesias R, Dahl G, Qiu F, Spray DC and Scemes E. Pannexin 1: the molecular substrate of astrocyte "hemichannels". J Neurosci 29: 7092–7097, 2009.

74. Syeda R, Qiu Z, Dubin AE, Murthy SE, Florendo MN, Mason DE, Mathur J, Cahalan SM, Peters EC, Montal M and Patapoutian A. LRRC8 proteins form volume-regulated anion channels that sense ionic strength. Cell 164: 499–511, 2016. doi: 10.1016/j.cell.2015.12.031

75. Concepcion AR, Wagner LE, Zhu J, Tao AY, Yang J, Khodadadi-Jamayran A, Wang YH, Liu M, Rose RE, Jones DR, Coetzee WA, Yule DI and Feske S. The volume-regulated anion channel LRRC8C suppresses T cell function by regulating cyclic dinucleotide transport and STING-p53 signaling. Nat Immunol 23: 287–302, 2022. doi: 10.1038/s41590-021-01105-x

76. Yu Q, Zhao Y, Maurer J, Arullampalam P, John N, Tranter JD, Abd El-Aziz TM, Lin M, Lerner DJ, Halabi CM and Sah R. Endothelial LRRC8C associates with LRRC8A and LRRC8B to regulate vascular reactivity and blood pressure. bioRxiv 10.1101/2025.08.17.670763, 2025. doi: 10.1101/2025.08.17.670763

77. Groulx N, Boudreault F, Orlov SN and Grygorczyk R. Membrane reserves and hypotonic cell swelling. J Membr Biol 214: 43–56, 2006.

78. Li P, Hu M, Wang C, Feng X, Zhao Z, Yang Y, Sahoo N, Gu M, Yang Y, Xiao S, Sah R, Cover TL, Chou J, Geha R, Benavides F, Hume RI and Xu H. LRRC8 family proteins within lysosomes regulate cellular osmoregulation and enhance cell survival to multiple physiological stresses. Proc Natl Acad Sci U S A 117: 29155–29165, 2020.

79. Kumar A, Zhao Y, Xie L, Chadda R, Tranter JD, Mikami RT, Abraham N, Hong J, Feng E, Rawnsley DR, Liu H, Henry KM, Meyer G, Hu M, Xu H, Hinton A, Jr., Grueter CE, Abel ED, Norris AW, Diwan A and Sah R. Lysosomal LRRC8 complex impacts lysosomal pH, morphology, and systemic glucose metabolism. Sci Adv 11: eadt6366, 2025. doi: 10.1126/sciadv.adt6366

80. Ghosh A, Khandelwal N, Kumar A and Bera AK. Leucine-rich repeat-containing 8B protein is associated with the endoplasmic reticulum Ca(2+) leak in HEK293 cells. J Cell Sci 130: 3818–3828, 2017.

81. Yanushkevich S, Zieminska A, Gonzalez J, Anazco F, Song R, Arias-Cavieres A, Granados ST, Zou J, Rao Y and Concepcion AR. Recent advances in the structure, function and regulation of the volume-regulated anion channels and their role in immunity. J Physiol 2024. doi: 10.1113/JP285200

82. Quinodoz M, Rutz S, Peter V, Garavelli L, Innes AM, Lehmann EF, Kellenberger S, Peng Z, Barone A, Campos-Xavier B, Unger S, Rivolta C, Dutzler R and Superti-Furga A. De novo variants in LRRC8C resulting in constitutive channel activation cause a human multisystem disorder. EMBO J 44: 413–436, 2025. doi: 10.1038/s44318-024-00322-y

83. Ridder MC, Boor I, Lodder JC, Postma NL, Capdevila-Nortes X, Duarri A, Brussaard AB, Estevez R, Scheper GC, Mansvelder HD and van der Knaap MS. Megalencephalic leucoencephalopathy with cysts: defect in chloride currents and cell volume regulation. Brain 134: 3342–3354, 2011.

84. Capdevila-Nortes X, Lopez-Hernandez T, Apaja PM, Lopez de HM, Sirisi S, Callejo G, Arnedo T, Nunes V, Lukacs GL, Gasull X and Estevez R. Insights into MLC pathogenesis: GlialCAM is an MLC1 chaperone required for proper activation of volume-regulated anion currents. Hum Mol Genet 22: 4405–4416, 2013.

85. Dubey M, Brouwers E, Hamilton EMC, Stiedl O, Bugiani M, Koch H, Kole MHP, Boschert U, Wykes RC, Mansvelder HD, van der Knaap MS and Min R. Seizures and disturbed brain potassium dynamics in the leukodystrophy megalencephalic leukoencephalopathy with subcortical cysts. Ann Neurol 83: 636–649, 2018.

